# HP1a promotes chromatin liquidity and drives spontaneous heterochromatin compartmentalization

**DOI:** 10.1101/2024.10.18.618981

**Authors:** Lucy Brennan, Hyeong-Ku Kim, Serafin Colmenares, Tatum Ego, Je-Kyung Ryu, Gary Karpen

## Abstract

Compartmentalization of the nucleus into heterochromatin and euchromatin is highly conserved across eukaryotes. Constitutive heterochromatin (C-Het) constitutes a liquid-like condensate that packages the repetitive regions of the genome through the enrichment of histone modification H3K9me3 and recruitment of its cognate reader protein Heterochromatin Protein-1 (HP1a). The ability for well-ordered nucleosome arrays and HP1a to independently form biomolecular condensates suggests that the emergent material properties of C-Het compartments may contribute to its functions such as force-buffering, dosage-dependent gene silencing, and selective permeability. Using an *in vitro* reconstitution system we directly assess the contributions of H3K9me3 and HP1a on the biophysical properties of C-Het. In the presence of HP1a, H3K9me3 (Me-) and unmodified (U-) chromatin form co-condensates composed of distinct, immiscible domains. These chromatin domains form spontaneously and are reversible. Independently of HP1a, H3K9me3 modifications are sufficient to increase linker-DNA length within chromatin arrays and slow chromatin condensate growth. HP1a increases the liquidity of chromatin condensates while dramatically differentiating the viscoelastic properties of Me-chromatin versus U-chromatin. Mutating key residues in HP1a show that HP1a interactions with itself and chromatin determine the relative interfacial tension between chromatin compartments, however the formation of condensates is driven by the underlying chromatin. These direct measurements map the energetic landscape that determines C-Het compartmentalization, demonstrating that nuclear compartmentalization is a spontaneous and energetically favorable process in which HP1a plays a critical role in establishing a hierarchy of affinities between H3K9me3-chromatin and unmodified-chromatin.

**Highlights:**

⍰ HP1a is necessary and sufficient for heterochromatin compartmentalization.
⍰ Heterochromatin compartmentalization is reversible and proceeds through microphase-separation.
⍰ H3K9me3 is sufficient to change nucleosome-array dynamics and mesoscale material properties.
⍰ HP1a increases chromatin liquidity.
⍰ HP1a-chromatin interaction modes tune the interfacial tensions and morphologies of heterochromatin compartments.

## Introduction

Heterochromatin and euchromatin are the two major chromatin compartments within the nucleus. Constitutive Heterochromatin (C-Het) is generally defined as telomere and centromere-proximal genome regions enriched in repetitive DNA sequences, H3K9me3-modified nucleosomes, and the cognate reader protein Heterochromatin Protein-1 (HP1a in *Drosophila*). C-Het plays a critical role in genome stability by repressing recombination within repetitive DNA, silencing expression and mobilization of transposable elements, and acting as a physical scaffold for the assembly of the kinetochore and sister cohesion^1^. Once formed, C-Het plays a global role in buffering the genome from both external and internal forces and contributes to the overall viscoelastic character of the nucleus^2,3^. Loss of HP1a or defects in H3K9me3 deposition leads to loss of transcriptional silencing, mitotic failures, loss of nuclear compartmentalization, and large-scale defects in nuclear morphology^2,4,5^. Smaller (10’s of Kbp) islands of heterochromatin located within otherwise euchromatic chromosome arms function to repress specific genes to promote cellular differentiation and maintain cellular identity^6^. Despite the large linear separation of C-Het and heterochromatin islands along the chromosomes, these smaller heterochromatin regions spatially associate with larger constitutive heterochromatin bodies^7,8^. The self-associative property of heterochromatin is most apparent at the initial stages of C-het growth during early embryo development, where small puncta of H3K9me3 and HP1a undergo large-scale reorganization and growth, both linearly along the chromosome^8^ and in three dimensions via liquid-like fusions^9,10^. The liquid-like dynamics of C-het *in vivo,* together with the *in vitro* findings that HP1 proteins undergo phase-separation to form biomolecular condensates, has prompted the hypothesis that C-Het is a biomolecular condensate that forms via phase-separation within the nucleus^9,11–13^.

*In vitro*, the formation of liquid-like condensates by either chromatin or (unphosphorylated)HP1a both require an underlying polymer: chromatin array under four nucleosomes do not phase-separate, nor does HP1a in the absence of DNA, RNA, or chromatin^11,14–16^. Polymer-based phase separation systems fall under the definition of complex coacervation, where charged polyelectrolytes spontaneously demix from their solvent^17,18^. *In vivo,* there are many examples of complex coacervation and liquid-liquid phase separation in the nucleus, for example the nucleolus and Cajal bodies^19–21^. The diversity in biocondensate composition and interaction hierarchies directly determines the biophysical properties of these compartments, such as their growth dynamics, viscosity, and interfacial tensions. How to accurately measure the biophysical properties of chromatin-based biomolecular condensates *in vivo* has presented a unique challenge due to the large range of length scales over which the chromatin functions^22,23^. C-Het represents an extreme example of this as it spans three orders of magnitude in length scales. A single H3K9me3 mark is on the order of Angstroms^24^, which can be bound by 2-4 HP1a molecules at the nanometer scale^16,25^, and these HP1a-coated H3K9me3 domains spread along megabases of linear DNA forming 3-dimentional structures on the order of micrometers^26^. Furthermore, changes in the molecular composition of C-Het compartments over the lifetime of an organism and across cell-types, adds an additional layer of complexity related to how biomolecular condensates, particularly those containing polymers, age^27,28^. Thus, it’s no wonder that there is confusion about the liquid-like nature of heterochromatin and the underlying biophysical principles that describe the formation and material properties of C-Het condensates^29–31^.

Currently, the biophysical properties of heterochromatin components have been inferred based on separate studies of either HP1a or chromatin. Our current understanding of the multi-dimensional network of HP1 interactions, and their impact on *in vivo* functions, are primarily derived from studies of HP1a (Su(var)205) in Drosophila, HP1α (Cbx5) in mammals, and Swi6 in *S. pombe* (hereafter referred to as HP1a, reviewed in ^32^). HP1a proteins contain multiple domains with distinct binding affinities: a disordered N-terminus extension, the structured H3K9me3-binding chromodomain (CD), a disordered hinge region, the dimerizing chromoshadow domain (CSD), and a disordered C-terminal extension. The CD contains the H3K9me3 recognition cleft and is required for heterochromatin localization of HP1a^33–37^.

Dimerization via the CSD is also required for heterochromatin association^2,38–40^ by promoting conformational changes within HP1^31^, direct interaction between the CSD and histone H3^41,42^, binding of HP1 partner proteins, and coordination with the CD^43^. The charged hinge region interacts with nucleic acids, chromatin, and promotes HP1a-oligomerization^15,16,44^. Cooperativity between all these domains has been implicated in the observed increase in affinity and specificity HP1a has for H3K9me3-nucleosome arrays as compared to mono-nucleosomes or H3K9me3-peptides^16,43,45,46^. The hinge region and CSD have also been shown to be essential for HP1a phase-separation, albeit at non-physiological salt conditions^11,15,16^.

Separately, it has been shown that well-ordered chromatin arrays can independently form liquid-like condensates at physiological salt conditions^14,47,48^. Such arrays are reminiscent of the nucleosome ‘clutches’ observed within the interphase nucleus ^49^. The inherent ability of chromatin to self-associate and form condensates is modulated by the length of linker DNA and disrupted by histone acetylation, and the material properties can be modulated by incorporation of peptides and linker histone H1^14,45,50–52^. Molecular dynamics simulations have provided a coarse-grained picture of how nucleosome spacing and histone tails modulate the flexibility of the chromatin polymer and facilitate a transient and diverse network of chromatin interactions in cis- and trans-, which are required for condensate formation^50,53–55^. Histone tails, specifically of H3 and H4, play a critical role in nucleosome-nucleosome interactions^51,56^ and are essential for chromatin condensate formation^14^. Importantly, the majority of post-translational modifications used to define nuclear compartments are deposited on the H3 and H4 tails^57,58^. Structural data shows that specific modifications change the balance of intra- and inter-nucleosome interactions and may thus modulate chromatin condensate properties in unique ways^14,45,59^. However, to date the only histone PTM characterized in the context of chromatin condensates is non-specific acetylation, which results in the dissolution of pre-formed condensates^14^.

The first step towards understanding how HP1a and H3K9me3-chromatin synergize to define heterochromatin compartments in the nucleus is to quantify the individual effects of H3K9me3 and HP1a on chromatin condensate formation and their emergent biophysical properties. To do this we have implemented an *in vitro* reconstitution system that leverages the ability of both HP1a and chromatin to form liquid-like condensates. We find that H3K9me3 impacts both the mesoscopic and nanoscopic properties of chromatin condensates but is not sufficient to drive macroscopic compartmentalization of unmodified (U-) chromatin from H3K9me3-(Me-) chromatin *in vitro.* HP1a is necessary for the formation of distinct Me- and U-chromatin compartments, and the degree of chromatin compartmentalization is determined by both HP1a-HP1a and HP1a-Me-chromatin interactions. This *in vitro* system provides a direct measure of the energetic landscape that determines C-Het compartmentalization, demonstrating that nuclear compartmentalization is a spontaneous and energetically favorable process in which HP1a plays a critical role in establishing a hierarchy of affinities between H3K9me3-chromatin and unmodified-chromatin.

## Results

### HP1a drives the formation of distinct chromatin compartments

To determine the minimal requirements the formation of C-het compartments, we took advantage of the ability for well-ordered chromatin arrays to form condensates^14^ to test weather H3K9me3 and/or HP1a are capable of driving chromatin compartmentalization. Reconstituted 12-nucleosome arrays containing either unmodified nucleosomes (U-chromatin) or H3K9me3-nucleosomes (Me-chromatin) were mixed with recombinant HP1a at physiological salt conditions (Figure 1A). In the absence of HP1a, U-chromatin and Me-chromatin mix and form homogeneous co-condensates (Figure 1B). In contrast, the presence of HP1a results in demixing of U- and Me-chromatin, and enrichment into distinct macroscopic compartments containing little of the other chromatin-type (Figure 1B). The partitioning of each chromatin type within the droplets provides a metric to quantify the relative interaction strengths between Me- and U-chromatin, as well as the relative interactions between each chromatin type and the surrounding solvent. In the absence of HP1a, both Me- and U-chromatin have a partitioning coefficient of 1 within the condensates (Figure 1C), indicating that the relative “strength” of Me- and U-chromatin interactions is approximately equal to the interaction of either chromatin type with the surrounding solvent. This is significantly different than in the presence of HP1a, where on average, 75% of the total Me-chromatin within each droplet partitions into a distinct Me-compartment. The remaining 25% of Me-chromatin is found within the U-chromatin compartment, resulting in a partition coefficient of ∼3 (Figure 1C). This neatly corresponds to the enrichment of approximately 75% of the total HP1a within the Me-chromatin compartments, also resulting in an HP1a-partition coefficient of approximately 3 (Figure 1C). The compartmentalization within individual droplets indicates that HP1a imparts an asymmetry to the interaction between Me- and U-chromatin. The chromatin compartments partially wet each other, and U-chromatin domains are more frequently observed to be completely coated by Me-chromatin indicating that U-chromatin-solvent interactions are less favorable than Me-chromatin-solvent interactions (SI figure 1, SI movies 1-4).

**Figure 1.**
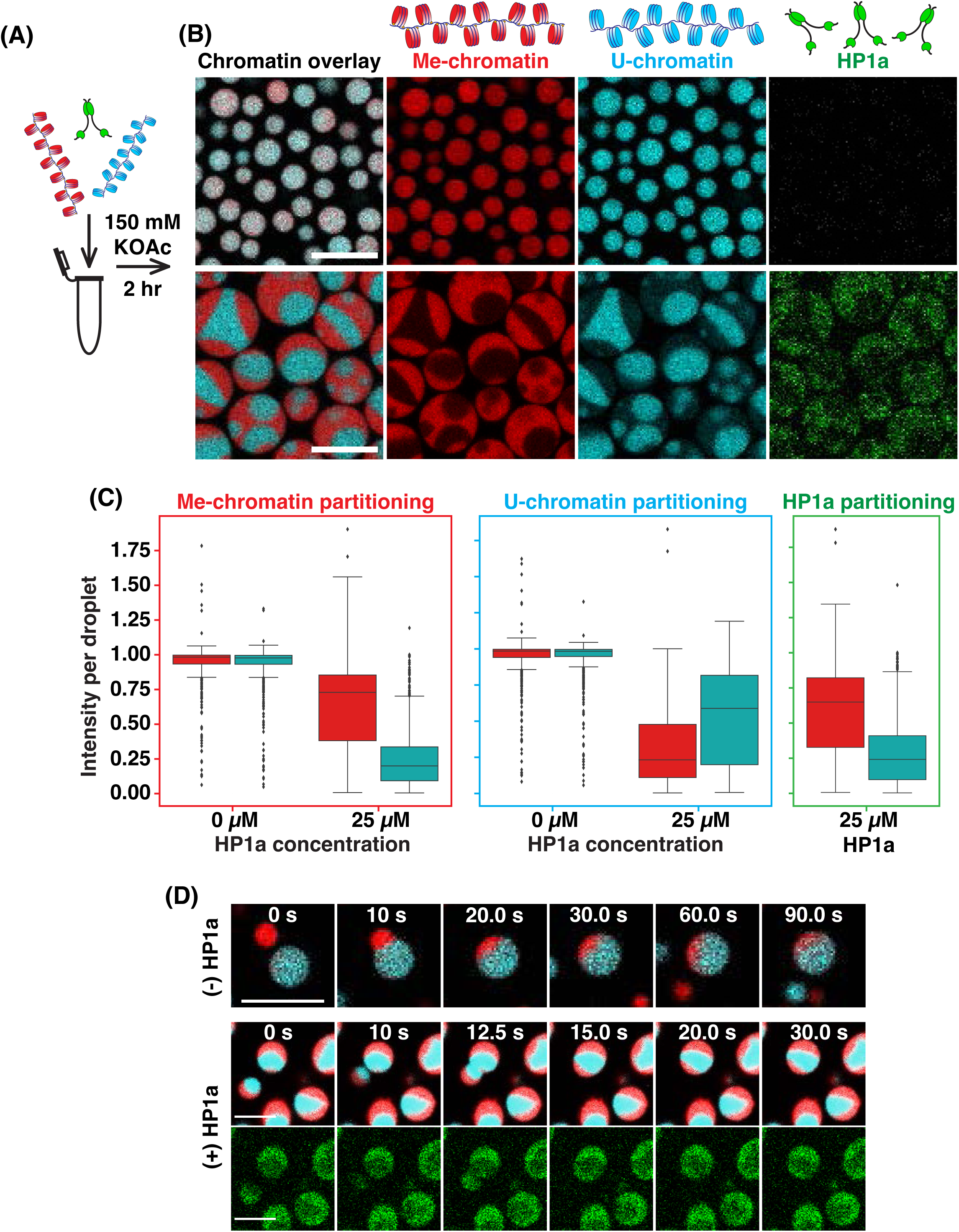
HP1a compartmentalizes Me- and U-chromatin under physiological salt concentrations. **(A)** Schematic of the experimental set-up, where the indicated components are mixed together in a physiological buffer, allowed to react for approximately 2 hours, then imaged. See methods for more details. **(B)** Representative images of Me- and U-chromatin mixtures (red and cyan respectively, top row) and Me-, U-chromatin, and HP1a mixtures (in green) (bottom row). Scale bar is 10 μm, overlay displays only the fluorescence signals of the chromatin. **(C)** Drop-wise partitioning of each reaction component (Me-chromatin, U-chromatin, or HP1a) into either Me-chromatin regions (red boxes) or U-chromatin region (cyan boxes) at 0 μμ and 25 μμ concentrations of HP1a. The boxes represent the quartiles of the data with the middle line being the median value of the distribution. Whiskers represent the full range of the data, outliers are shown as black diamonds. **(D)** representative fusions of mixed droplets in the absence (-HP1a) or presence (+HP1a) of HP1a. Scale bar is 5 μm in all images and the timescale is indicated.

This asymmetry in interactions also modulates the formation dynamics of chromatin condensate mixtures, as is predicted for a system undergoing spontaneous demixing from the surrounding solvent. Time-lapse imaging shows that in the absence of HP1a, both Me- and U-chromatin forms phase-separated condensates capable of fusing together and undergoing slow internal mixing (Figure 1D, 0 M HP1a, SI Figure 2A). The internal chromatin compartments formed in the presence of HP1a are observed from the outset of condensate formation, are stable throughout fusion events, and display a preference for fusions to take place between U-compartments (Figure 1D, 25 M HP1a, SI Figure 2B). To determine if HP1a was retarding the mixing dynamics of the underlying chromatin-polymer, irrespective of histone modification state, mixtures of a single chromatin type (unmodified or H3K9me3), labeled with cy3 or cy5, were mixed with HP1a. These mixtures resulted in homogeneously mixed condensates (SI Figure 3). These data demonstrate that the formation of distinct compartments requires HP1a and two populations of chromatin harboring different densities of H3K9me3.

**Figure 2.**
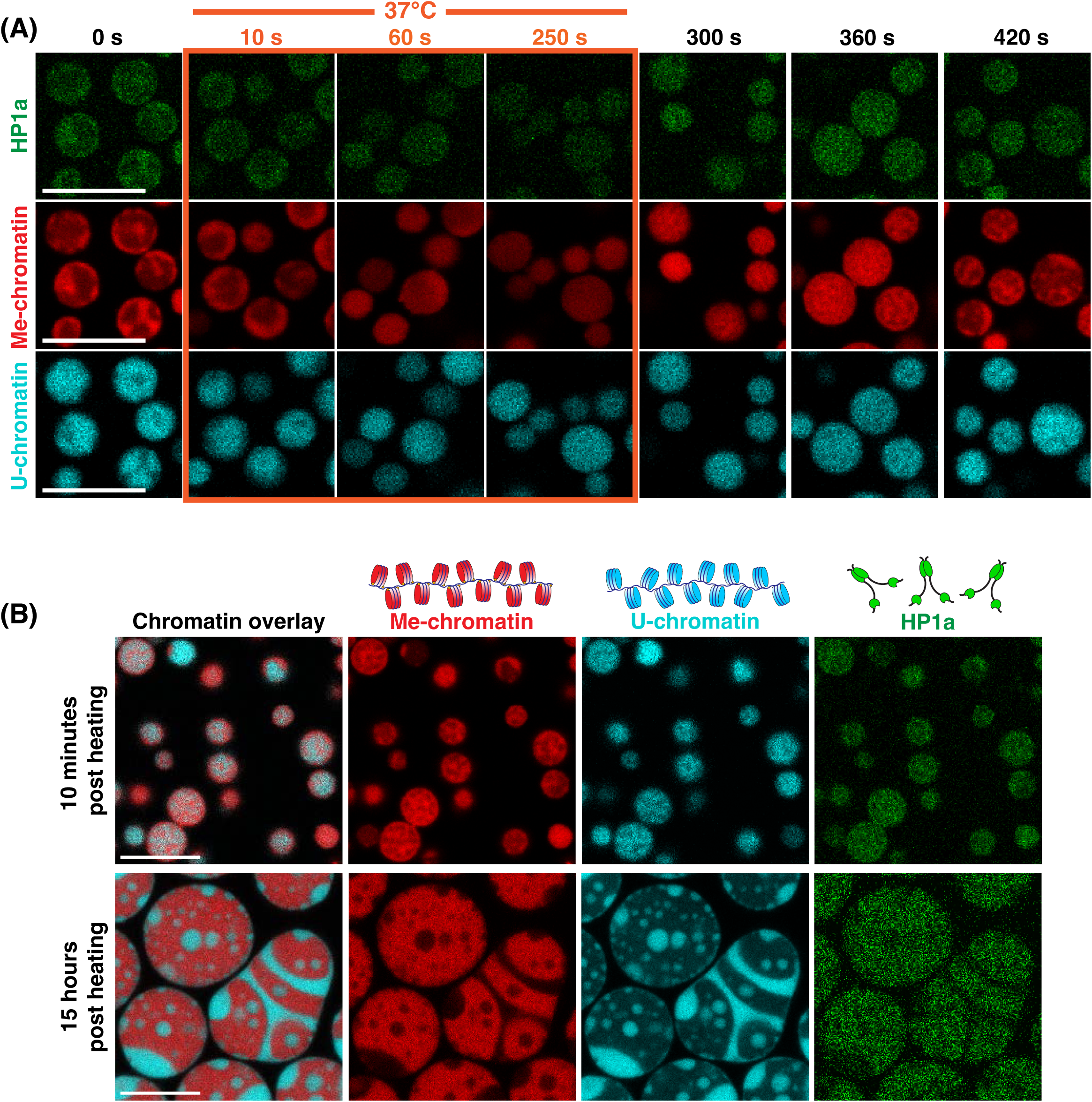
HP1a-chromatin compartments are reversible. **(A)** Time-course of droplet heating experiments demonstrating loss of compartments and mixing of U- and Me-chromatin upon heating, plus microphase demixing upon cooling. Each frame shows different droplets due to the convective flows generated by the heating device. **(B)** Representative droplets from early (10 min) and late (15 hrs) time points post heating demonstrate demixing and reformation of distinct compartments. Scale bar is 10 μm in all images.

**Figure 3.**
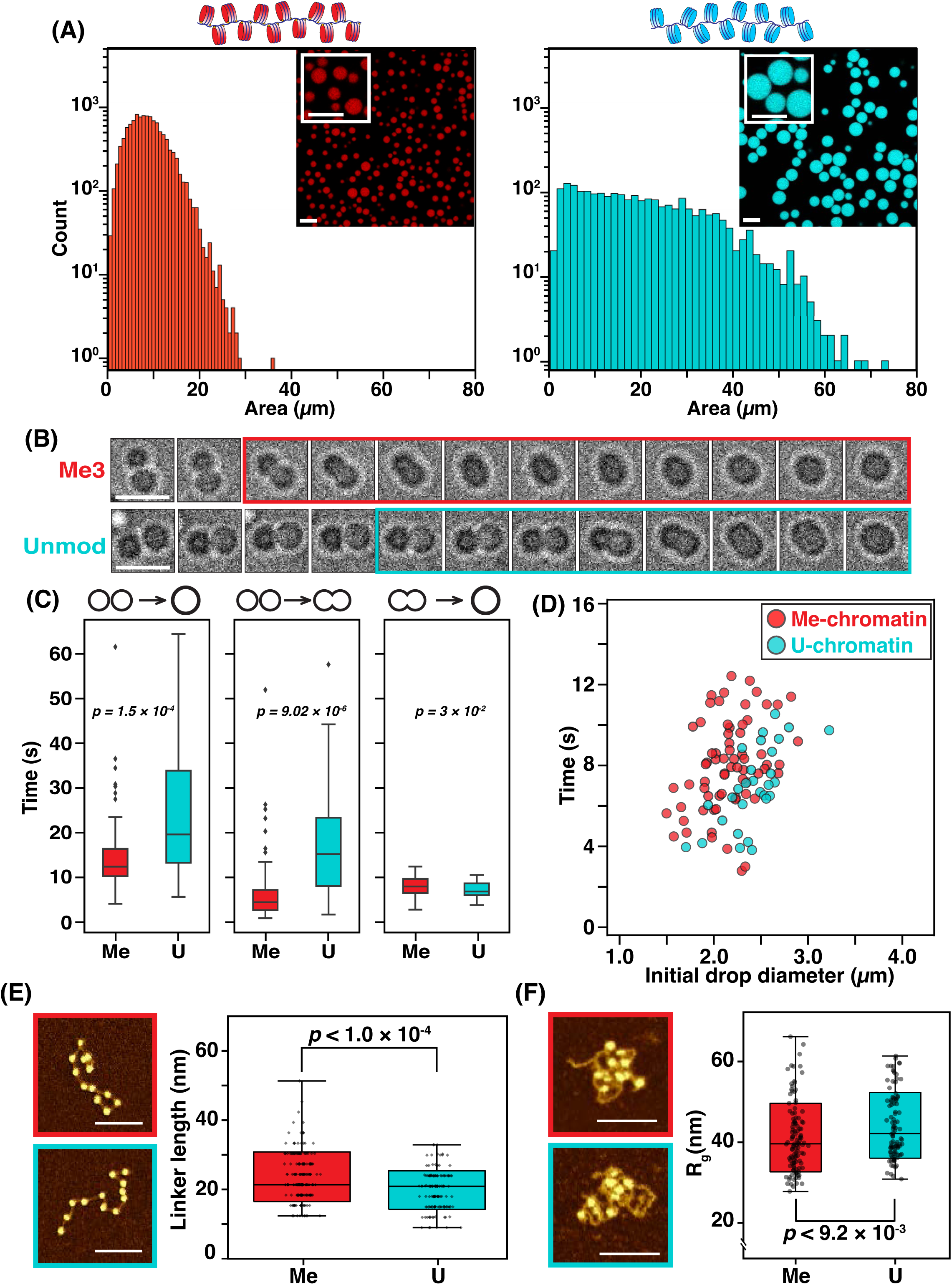
H3K9me3 is sufficient to change chromatin material properties. **(A)** Histogram of droplet areas (in μm) for Me-chromatin condensates (red, left) and U-chromatin condensates (cyan, right). Images are full fields of view and insets are zoomed images (scale bars are 5 μm). **(B)** Representative brightfield time course of droplet fusions for Me-condensate (top) and U-condensates (bottom). Boxes indicate the time plotted in panel D. Each frame is consecutive with a 1 second interval. Starting frame represents the “end” of time-1, after stable droplet contact has been observed but before the onset of time-2. Scale bar is 5 μm. **(C)** Boxplot distributions for three fusion timescales (see cartoon diagram and main text). The boxes represent the quartiles of the data with the middle line being the median value of the distribution. Whiskers represent the full range of the data, outliers are shown as black diamonds. **(D)** Full distribution of the second fusion time in seconds versus average initial droplet diameter in μm. Red is data for Me-condensates and cyan is for U-condensates. **(E)** Representative AFM images of individual chromatin arrays used for linker-length analysis; red box is Me-chromatin, cyan box is U-chromatin. Box plot displays the distribution of linker lengths measured for Me-chromatin arrays (red) or U-chromatin arrays (cyan). All data points overlaid in black. **(F)** Representative AFM images of individual chromatin arrays used for R_g_ analysis; red box is Me-chromatin, cyan box is U-chromatin. Box plot displays the distribution of radius of gyration measured for Me-chromatin arrays (red) or U-chromatin arrays (cyan). All data points overlaid in black. Scale bars for all AFM images are 100 nm.

The preferential self-association of Me- and U-chromatin requires HP1a and results in distinct Me- and U-chromatin domains that adhere to one another creating highly deformable yet stable internal interfaces between the chromatin compartments reminiscent of those observed within the interphase nucleus. We conclude that HP1a drives the formation of chromatin compartments by modulating both Me- and U-chromatin interactions, as well as the interactions of chromatin with the surrounding solvent.

### Chromatin compartmentalization by HP1a is reversible and proceeds through micro-phase separation intermediates

The stability of Me- and U-chromatin domains in the presence of HP1a recalls the observation that mammalian HP1α (Cbx5) stabilizes distinct DNA domains in condensates formed under low-salt conditions^15^. This raises the possibility that the domains observed in the context of chromatin are also due to kinetic trapping of the underlying polymer by HP1a. If this is indeed the case, adding sufficient energy to disrupt HP1a cross-linking would allow the polymer to relax into a homogeneously mixed state, which would remain homogeneous upon removal of the external energy source. To test this hypothesis, heating experiments were performed to determine the energetic barrier separating Me- and U-chromatin domains within the tripartite mixtures.

Heating the tripartite condensates to 37°C is sufficient to disrupt HP1a-dependent compartmentalization of U- and Me-condensates (Figure 2A). This corresponds with the temperature required to dissociate HP1a from the chromocenter in cultured *Drosophila* S2 cells (SI Figure 4), indicating good correspondence between the energy landscape for HP1a binding between *in vivo* and *in vitro* systems. After incubating the *in vitro* system for one minute at 37°C, the chromatin and HP1a are uniformly distributed through the condensates (Figure 2A, SI Movie 7). Upon release of the heating set-point, the chamber cools to room temperature, and within 90 seconds, small Me-chromatin domains appear within the condensates. The microdomains grow and fuse over time, ultimately reverting to the macroscopic domains observed in our initial mixing experiments (Figure 2B, SI Movie 7). The internal demixing of chromatin types demonstrates that these distinct compartments form spontaneously and represent a true energetic minimum of the system, rather than a kinetically arrested intermediate state. Furthermore, these data demonstrate that HP1a is sufficient to drive the self-association of Me-chromatin into macroscopic domains within a dense chromatin environment, analogous to C-Het coalescence observed in the early *Drosophila* embryo^9,60^.

**Figure 4.**
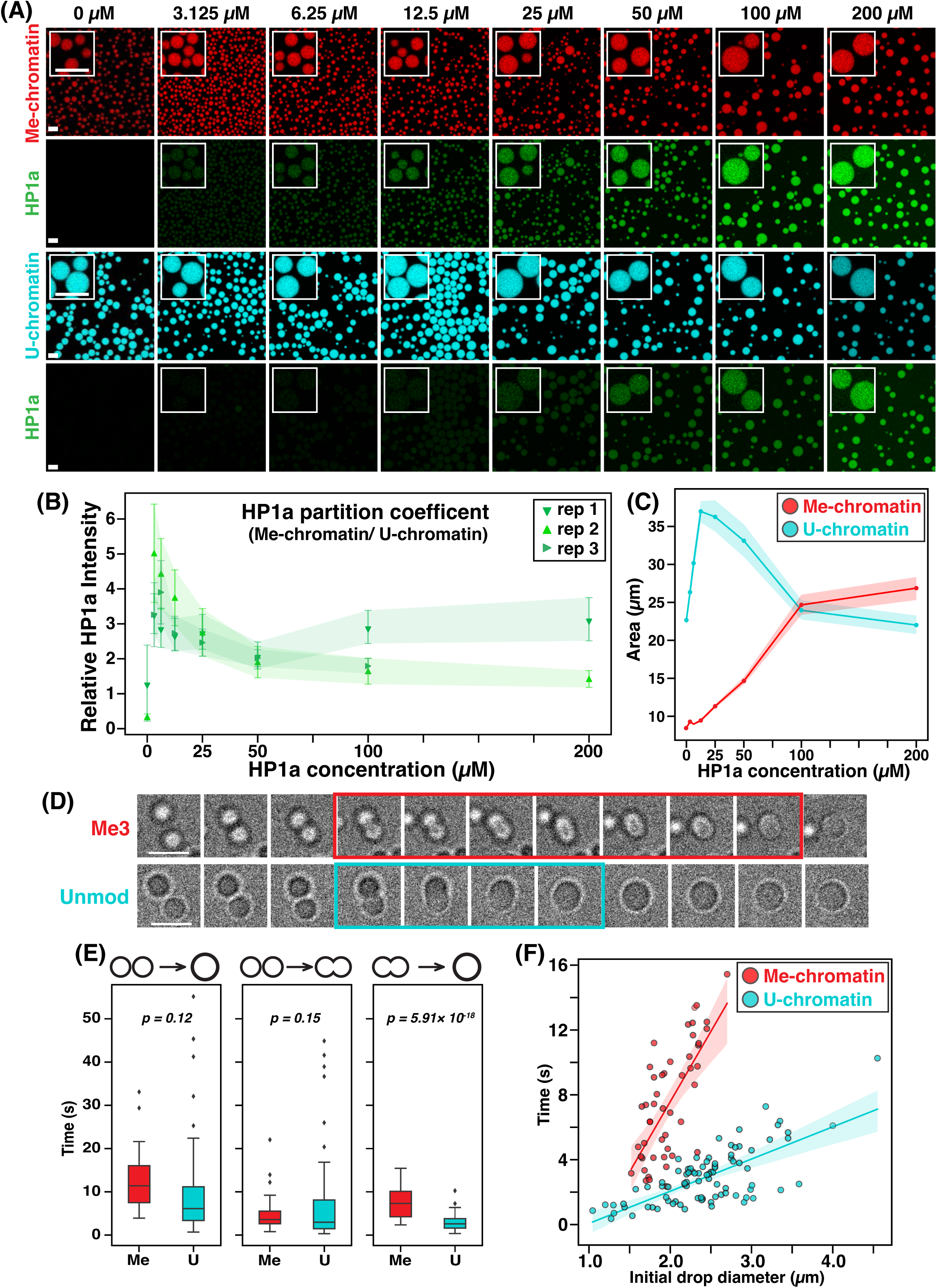
HP1a liquifies chromatin condensates. **(A)** HP1a titration into Me-condensates (red, top 2 rows) and U-condensates (cyan, bottom 2 rows). Images are full fields of view and insets are zoomed from the same image. Scale bars are 5 μm. **(B)** HP1a partition coefficient. Plot shows the relative HP1a-intensity in Me-condensates divided by HP1a-intensity in U-condensates at each HP1a concentration, for three separate replicates (shades of green). Filled triangles represent the average partition coefficient, bars/shading are the standard deviation. **(C)** Plot displaying the average (filled circle) chromatin condensate size at each HP1a concentration, shading shows 95% confidence interval (Me in red, U in cyan). **(D)** Representative brightfield time-courses of droplet fusions for Me-condensate (top) and U-condensates (bottom). Boxed frames indicate time-2 plotted in panel D. Each frame is consecutive with a 1 second interval. Starting frame represents time = 0. Scale bar is 5 μm. **(E)** Boxplot distributions for three fusion time components (see cartoon diagram and main text). The boxes represent the quartiles of the data with the middle line being the median value of the distribution. Whiskers represent the full range of the data, outliers are shown as black diamonds. **(F)** Full distribution of the second fusion time in seconds versus average initial droplet diameter in μm. Red is data for Me-condensates and cyan is for U-condensates.

### H3K9me3 modulates nano- and meso-scale nucleosome dynamics

To further dissect the effects of C-Het core components on the material properties of chromatin condensates, we further simplified the system to examine what, if any, effects H3K9me3 may have on chromatin condensate formation. Prior work has shown that nucleosome spacing and post-translational modification can promote (10n+5 spacing) or inhibit (global acetylation) chromatin condensate formation^14^. Here, U-nucleosomes and Me-nucleosomes were assembled on the same underlying DNA and therefore should have the same linker length.

However, in isolation, Me-chromatin formed smaller (mean=8.9 ± 4.13μm^2^), more monodispersed droplets, whereas U-chromatin formed larger (mean =22.8 ± 13.81μm^2^) condensates with a broad distribution of sizes (Figure 3A). These dramatically different distributions suggest that H3K9me3 may modulate condensate growth and/or the internal packing of Me-chromatin within the condensate.

Partitioning measurements of two sizes of fluorescent, anionic dextrans did not reveal any differences in Me- or U-chromatin condensate mesh sizes (SI figure 5), suggesting that the observed differences in droplet size are due to differential growth dynamics rather than chromatin packing. Cryo-ET studies have described the growth of chromatin condensates as occurring via two distinct stages; initial nucleation of irregular clusters and mesoscale growth via accretion and fusion^61^. The measurements of droplet size were performed at 2 hours following the initial mixing of all reaction components, a time at which the majority of droplets ceased fusing and settled on the surface of the well. This allowed for inferences about the growth dynamics of each system based on their size distribution at a given time after the onset of phase-separation on-set. The broad distribution of U-chromatin sizes (Figure 3A) suggests that these condensates primarily grow in a stochastic manner through many diffusion-driven collision events. The well-defined size distribution of Me-chromatin condensates indicates that they have undergone fewer fusion events than U-condensates, in the same amount of time.

**Figure 5.**
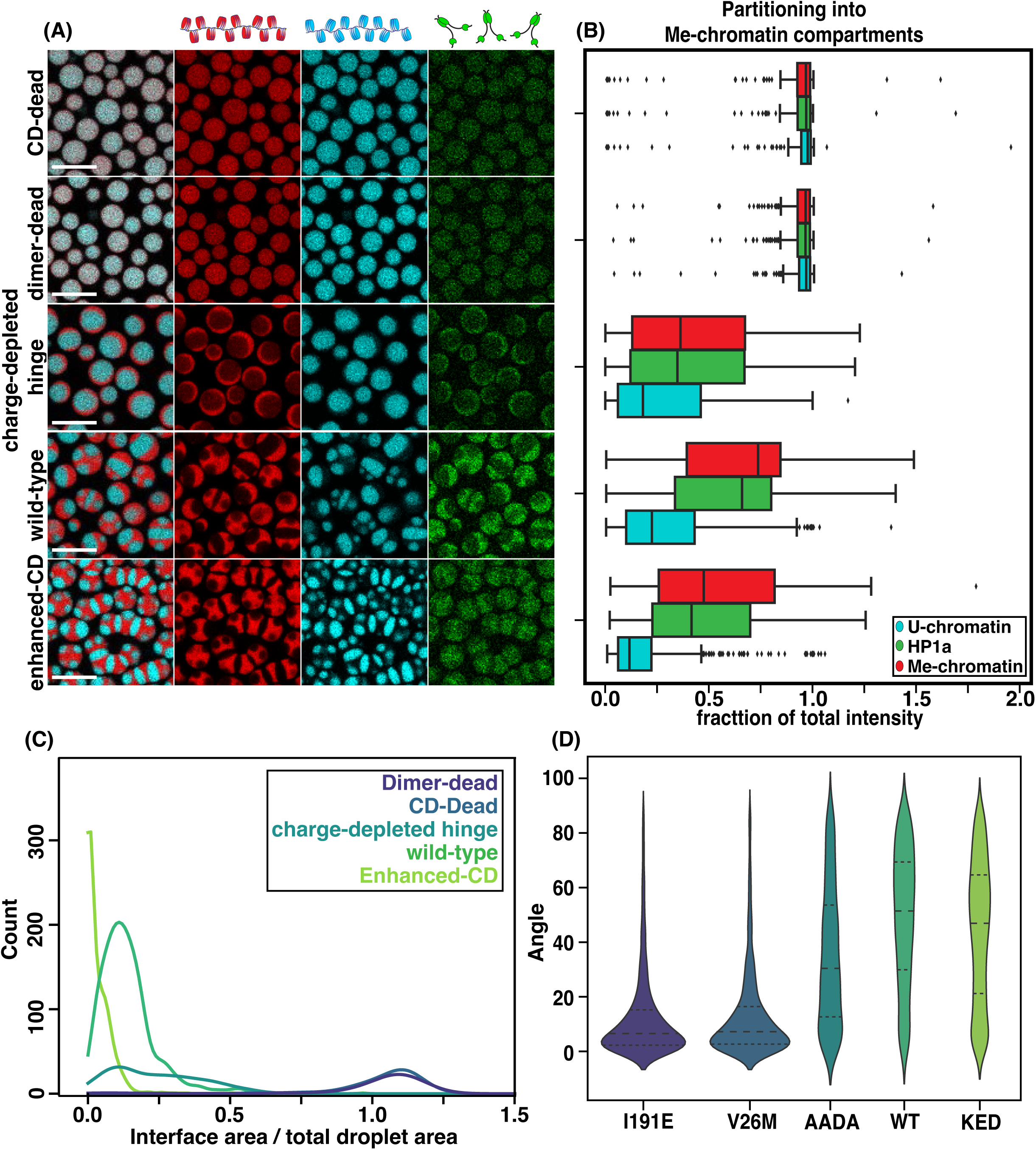
HP1a mutants modulate the interfacial tensions of chromatin condensates. **(A)** Representative images of Me-, U-chromatin, and HP1a-mutant mixtures (red, cyan, and green, respectively). Scale bar is 10 μm, overlay displays only the fluorescence signals of the chromatin (far left column). **(B)** Drop-wise partitioning of each condensate component into the Me-chromatin regions. The boxes represent the quartiles of the data with the middle line being the median value of the distribution. Whiskers represent the full range of the data, outliers are shown as black diamonds. **(C)** Kernal density plot of area occupied by intersecting Me- and U-chromatin domains in each droplet, color coded by HP1a-mutant type. **(D)** violin plot of the contact angles measured between Me- and U-chromatin compartments within the same droplet for each mutant mixture. The inner lines represent the quartiles of the data with the middle line being the median value of the distribution. The outer violin is the default kernel density estimate used by seaborn.

To characterize the fusion dynamics of Me- or U-condensates, we monitored individual droplet fusion events measuring several time-scales along with droplet sizes (Figure 3B). Three timescales for droplet fusion were defined: (1) stable surface contact to complete coalescence (total-time = time1 + time2), (2) stable contact to half-fused (time1), and (3) half-fused to fully coalesced (time2). All of these timescales are determined by both internal viscosity and interfacial tensions^62^, and time2 can be used to quantify the ratio between viscosity and interfacial tensions^63^. Interestingly, Me-condensates displayed a significantly faster time-1 compared to U-condensates (Me:14.96 ± 8.81 s, U: 24.62 ± 14.98 s, p=9.02×10^-6^), suggesting that establishment of a stable contact interface occurs faster for Me-condensates than U-condensates. In contrast, time-2 is slightly but significantly slower for Me-condensates (Me: 8.00 ± 2.25 s, U:6.94 ± 1.98 s, p=0.03), suggesting that the interfacial tension and/or viscosity of Me-condensates is slightly higher than U-chromatin. Plotting the distribution of time-2 versus the initial starting diameter of the droplets shows that these distributions are indeed distinct, with U-condensates slightly shifted to faster times and larger droplets. The distribution of Me-condensates does not display a linear relationship, as would be expected for a viscoelastic material^63^. However, the U-condensate distribution fits moderately well (R^2^=0.68) to a line with a slope of ∼4.5 μm/sec (SI Figure 6), indicative of a viscoelastic material. The difference in fusion dynamics between Me- and U-condensates indicates that H3K9me3 slows the internal coalescence of condensates, however the initial merging of two droplets occurs faster for Me-condensates than for U-condensates. For Me-condensates to remain small, as they do, fewer fusion events take place, suggesting that there is a significant energetic barrier preventing the initial stages of Me-condensate coalescence.

**Figure 6.**
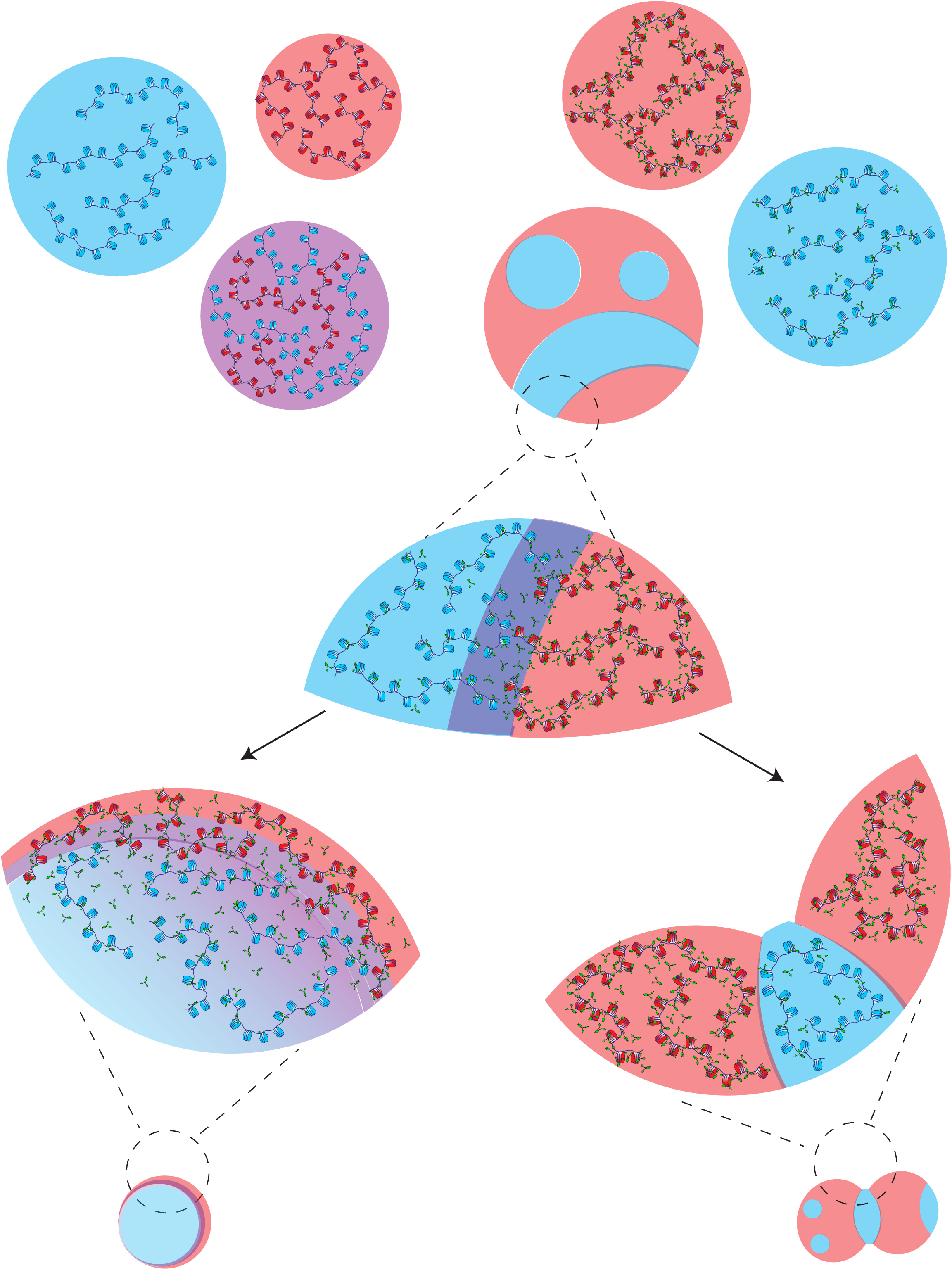
Model schematic of the Me- and U-chromatin interfaces formed in the presence of HP1a. In the absence of HP1a (top left) Me-chromatin condensates are smaller than U-condensates which we hypothesize is due to the decreased radius of gyration measured for Me-chromatin fibers. HP1a partitions more favorably into Me-chromatin condensates over U-chromatin condensates and increases the liquidity of both, resulting in larger droplets (top right). Mixtures of Me- and U-chromatin are homogenous in the absence of HP1a (purple droplet) but display clear compartmentalization with HP1a (red and blue droplet). The interface “strength” between Me- and U-chromatin compartments is governed by multimodal affinities of HP1a to the underlying chromatin (middle zoom). Mutations that decrease the positive charge of the HP1a hinge (KKRD ➔ AADA) weaken these interfaces, likely by weakening both HP1a self-association and HP1a-DNA affinity (bottom left). Mutations that increase HP1a affinity for H3K9me3 (KED) increase the segregation of Me- and U-chromatin reinforcing the interfaces between chromatin types (bottom right). In all tripartite mixtures, U-chromatin domains are consistently shielded from the surrounding solvent by Me-chromatin suggesting that HP1a differentially modulates chromatin solvation.

To determine what, if any, nanoscale effects H3K9me3 might have on the conformation of individual Me-chromatin arrays, we used atomic force microscopy to measure the length of linker DNA within individual arrays and assessed single-array compaction (SI Figure 7). There is a significant increase in linker-DNA length for the Me-chromatin array (mean: 23.28 nm) over U-chromatin (mean: 19.81 nm) corresponding to an average increase of 10 bp of linker DNA released by the H3K9me3 modification. Structurally, H3K9me3 sits right at the nucleosome dyad and the H3 tail has a substantial interaction footprint along the DNA exiting the nucleosome^64,65^. Our data indicates that H3K9me3 is sufficient to partially destabilize the wrapping of DNA as it exits the nucleosome (Figure 3E). To determine if this increase in linker-DNA by H3K9me3 changes the compaction of individual arrays, un-crosslinked arrays were deposited on an AFM grid and a proxy for radius of gyration was measured for single, well-separated chromatin arrays (SI Figure 8). Me-chromatin formed significantly more compact particles, suggesting that H3K9me3 favors more, or stronger, intra-array interactions (Figure 3E). This is consistent with previous work demonstrating that H3K9me3 chromatin arrays sediment faster than unmodified arrays ^45^.

Consistent with the hypothesis that H3K9me3 favors formation of intra-array contacts, chromatin droplets formed from individual arrays containing a random mixture of Me- and U-nucleosomes display a size distribution more akin to that measured for Me-saturated arrays (SI Figure 9). Interestingly, the size distributions of co-condensates formed by U-chromatin and Me-chromatin mixtures (Figure 1B) also displays a size distribution similar to that of Me-chromatin condensates, suggesting that H3K9me3 may modulate condensate size through both cis- and trans-interactions (SI figure 9).

Taken all together, we conclude that the liberation of linker DNA along with increased intra-array interactions limits fusion events of Me-condensates. Nucleosome arrays with linker lengths of 10n+5 (bp), display more disordered nucleosome conformations and an increase in long-range interactions compared to a 10n linker-length which favors intra-array interactions thus limiting inter-array interactions^50,54,55,66^. The extended conformation of the 10n+5 arrays favor inter-array interactions between disordered histone-tails and nucleosomes, and possibly DNA, on neighboring arrays. We suggest the increase in average linker length for Me-chromatin shifts nucleosome spacing away from the condensate-favoring 10n+5 bp, and thus limits the inter-array interactions necessary for droplet adhesion and internal rearrangements.

These findings suggest that while H3K9me3, independent of HP1a, is not sufficient for chromatin compartmentalization, this histone modification impacts Me-chromatin material properties and fusion dynamics by shifting the balance between intra- and inter-chromatin array interactions. This has implications in the context of the nucleus where clutches of nucleosomes saturated with H3K9me3 modifications may not interact as dynamically with adjacent nucleosome clutches. This bias towards more compact arrays is also reminiscent of the more compact morphology observed within C-Het domains; however our data suggests that this compaction is not due to closer linear packing of nucleosomes, but rather 3-dimensional compaction due to increased intra-array interactions.

### HP1a liquifies chromatin condensates

Given that H3K9me3 imparts a subtle, but significant, change in both the meso- and nanoscale properties of chromatin condensates (Figure 3), and HP1a modulates both U- and Me-chromatin solvation (Figure 1); we next asked how HP1a affects the material properties inherent to Me- and U-chromatin condensates. The ability of HP1 proteins (HP1a, HP1α, Swi6) to phase separate *in vitro* has been characterized both for the protein alone and in the presence of various substrates, though these studies were performed at non-physiological salt conditions^9,11,15,16,36^.

While providing valuable information about the mechanisms of HP1 self-association, these buffer conditions do not accurately recapitulate the interplay between the homotypic (HP1-HP1 or chromatin-chromatin) and heterotypic (HP1-chromatin) interactions that exist at *in vivo* ionic conditions.

To explore the effects of HP1a on chromatin condensates, we first sought to define a critical concentration for HP1a enrichment into either Me- or U-condensates. The concentration of HP1a was titrated into the system while nucleosome concentration was held constant (Figure 4A). At physiological salt conditions (150 mM monovalent salts), HP1a visibly enriched into Me-condensates across all tested concentrations, whereas enrichment into U-condensates was not evident at concentrations of HP1a below 12.5 μM (Figure 4A). The enrichment of HP1a into Me-condensates relative to U-condensates is most pronounced at the lower concentrations of HP1a and is never equally partitioned, even at high concentrations of HP1a (Figure 4B). There is satisfying agreement between the concentration at which HP1 enriches into chromatin condensates and the critical concentration at which HP1a drives chromatin phase-separation under low salt-conditions (75 mM K^+^). At these sub-physiological concentrations, chromatin does not phase separate in the absence of HP1a, however co-condensates of HP1a and chromatin form at 3.125μM HP1a with Me-chromatin and 25 μM HP1a with U-chromatin (SI Figure 10). This indicates that higher-order HP1a-HP1a interactions, which contribute to HP1a enrichment at physiological salt and HP1a-dependent phase-separation at low-salt, occur at a universal critical concentration. The effect of these HP1a-HP1a interactions on Me-condensates can be observed in the linear increase in droplet size with increasing HP1a concentration, suggesting that HP1a increases the number of fusion events over a given time (2 hr), leading to larger droplets (Figure 4C).

To probe the effects of HP1a enrichment on the material properties of chromatin condensates, individual fusion events were tracked over time. Across each timescale defined previously, U-condensates fuse faster than Me-condensates (Figure 4D, E). Particularly striking is the effect of HP1a on the relationship between time-2 and droplet size. In the presence of HP1a, both chromatin condensates now behave as viscoelastic materials; each with a distinct linear relationship between coalescence time (time-2) and droplet diameter (methyl: R^2^= 0.68, slope =8.66 μm/sec, unmod: R^2^= 0.71, slope: =1.98 μm/sec). This indicates that addition of HP1a increases the overall liquidity of chromatin, thus enhancing the ability of both U- and Me-chromatin condensates to undergo fusions. Interestingly, the ratio of viscosity to interfacial tensions (i.e. the reported slope) is significantly lower for U-chromatin condensates compared to Me-condensates, which could explain how heterochromatin acts to force-buffer the nucleus^2,67,68^.

### The interfacial tensions of chromatin domains are determined by HP1a-chromatin affinities

The viscoelastic properties of Me-chromatin versus U-chromatin are dictated by the fundamental interactions that govern chromatin phase-separation: Me-chromatin self-association, U-chromatin self-association, Me-chromatin-solvent interactions, U-chromatin-solvent interactions, and, when mixed, Me- and U-chromatin interactions. In a system of chromatin mixtures, these relative strengths of interactions can be measured by examining (1) the partitioning of chromatin into distinct compartments, (2) the relative amount of solvent vs. chromatin exposed interfaces, and (3) the contact angle between chromatin domains. The observation that both Me-chromatin and U-chromatin form spherical condensates is indicative of the unfavorable chromatin-solvent interactions which acts to minimize the surface area of solvent-exposed interfaces through the formation of phase-separated condensates. To dissect the effect of HP1a on Me- and U-chromatin interactions, four distinct HP1a mutants were purified and used in Me- and U-chromatin mixtures. The mutations disrupt one of the four main HP1a-chromatin interaction modes: the chromodomain (CD)-dead mutation (V26M ^35^) abolishes the interaction of HP1a with H3K9me3, the dimerization (CSD)-dead mutation (I191E ^69^) eliminates HP1a-HP1a dimerization, a charge-depleted hinge (AADA, Unpublished data S. Colmenares) disrupts HP1a-DNA interactions, and a CD-enhanced mutant (KED ^70^) decreases the dissociation of HP1a from H3K9me3 binding sites. When expressed in S2 cells, each of these mutants displays aberrant heterochromatin localization and distinct FRAP recovery curves representative of their relative affinity for Me-chromatin (SI Figure 11)

Chromatin compartmentalization by these HP1a mutants were compared with wild-type HP1a (Figure 5A). For wild-type HP1a, ∼75% of Me-chromatin and HP1a, per-droplet, partitions into a distinct compartment, in which ∼ 25% of the total U-chromatin also partitions (Figure 5B). Overall, wt-HP1a droplets are close to spherical and the Me-compartments tend to be located at the exterior, coating the U-chromatin domains. The area of overlap between Me-domains and U-domains occupies approximately 10-15% of the total droplet area (Figure 5C). Nearly 100% of the U-chromatin surface area (perimeter in 2-D) is found at a shared interface, compared to about 85% of the Me-chromatin interfaces (SI Figure 12B). The propensity for the U-chromatin to be engulfed by Me-chromatin, in the presence of HP1a, demonstrates that HP1a contributes to a higher interfacial tension (γ) between U-chromatin and the solvent relative to Me-chromatin and the solvent^23,71^. At the subset of interfaces where U-chromatin is exposed to the solvent, e.g. partially wet by Me-chromatin, the contact angle is widely distributed between 30°-70° (Figure 5D). The observation of partial wetting between U- and Me-chromatin compartments indicates that the interfacial tension between Me-and U-chromatin is higher than that of Me- or U-chromatin and the surrounding solvent. Thus, we can create a hierarchy of interfacial tensions: γ_me-u_ > γ_u-sol_ > γ_me-sol_, mediated by the interaction of HP1a with the underlying chromatin.

This is in stark contrast with the homogenously mixed droplets produced by either the CD-dead or dimerization-dead mutants (Figure 5A-C). In terms of the hierarchy of interfacial tensions, Me-U-chromatin interaction parameters for these mutant proteins are equal to that of chromatin and the solvent; γ_me-u_ = γ_u-sol_ = γ_me-sol_. The hierarchy of interfacial tensions established by wt-HP1 is partially rescued by the charge-depleted hinge mutant. This mutation converts a positive patch in the disordered hinge region to a neutral patch (KKDR ➔ AADA), reducing the net charge of the hinge from +4 to +1 (hinge pI =9.36 ➔ pI = 8.27). This mutant retains both dimerization and H3K9me3-recognition capabilities but forms domains of Me-chromatin that appear as thin layers coating the surface of U-chromatin domains (Figure 5A, SI Figure 13A).

The domains lack well-defined interfaces as evidenced by the decreased chromatin partitioning, increase in overlap between domains, and contact angles broadly distributed at about 30° (Figure 5). The Me-chromatin domains are exclusively observed at the surface of the droplets, acting to reduce the amount of solvent-exposed U-chromatin, supporting the hypothesis that U-chromatin-solvent interactions are more energetically costly than Me-chromatin-solvent interactions (Figure 5C, SI Figure 11A). At low salt, the charge-depleted hinge mutant and the dimerization-dead mutants are unable to form droplets with either Me- or U-chromatin (SI Figure 14 B,C). This indicates that both the self-association of HP1a along with the recruitment of HP1a to Me-chromatin is necessary to establish chromatin compartmentalization.

The KED mutant maintains all components of the HP1a interaction network, but increases the affinity between HP1a and H3K9me3 via three point mutations to the chromodomain binding site^70^. Chromatin mixtures containing the CD-enhanced HP1a are much more elongated in morphology and display increased chromatin compartmentalization and a decrease in overlap between chromatin compartments (Figure 5 A-C, SI Figure 11A). These droplets have a reduction in their shared surface area between U- and Me-chromatin domains, manifested as a significant decrease in U-chromatin domains embedded within Me-domains (SI Figure 12B, SI Figure 13B). However, the relative amount of U-chromatin exposed to the solvent is only mildly increased compared to wild-type. Instead, the U-chromatin domains become sandwiched between larger, more spherical Me-chromatin domains, and the amount of Me-compartment surface found at an interface decreases (Figure 5C, SI Figure 12B). Furthermore, the distribution of contact angles between the two chromatin compartments is slightly decreased from that observed for wt-HP1a (Figure 5D). These data suggest that the chromo-enhanced mutant selectively enhances the Me-chromatin self-interactions, which decreases the amount of shared interfaces between Me- and U-chromatin compartments.

Together, these data demonstrate that tuning HP1a-chromatin interaction modes alters the hierarchy of interfacial tensions that determine the integrity and morphology of chromatin compartments.

## Discussion

This study defines a minimal system capable of recapitulating heterochromatin compartmentalization with just three components: HP1a, unmodified, and H3K9-methylated nucleosome arrays. Utilizing this minimal system, we have quantified the molecular interactions that underlie constitutive heterochromatin compartmentalization in terms of (1) the energetics that give rise to co-existing chromatin compartmentalization and (2) the material properties of the resulting condensate(s). The compartmentalization of Me-chromatin away from U-chromatin by HP1a arises spontaneously within this tripartite system. Me-chromatin and U-chromatin domains associate, creating interfaces which can be tuned by modulating the HP1a and chromatin interaction affinities (Figure 6). The two chromatin compartments have distinct material properties, conferred by H3K9me3 modifications, and further differentiated by HP1a binding. HP1a increases chromatin liquidity of both U- and Me-chromatin while establishing a clear hierarchy of interfacial tensions; γ_u-sol_ ≥ γ_me-u_ > γ_me-sol_.

### Heterochromatin compartmentalization is a spontaneous and energetically favorable process

Distinct Me-chromatin and U-chromatin compartments arise only when these two chromatin-types are mixed in the presence of HP1a (Figure 1, Figure 6). No other components are required and these immiscible domains are observed throughout condensate growth, form spontaneously at the outset of visible condensate formation, and are stable through droplet fusion events. Importantly, these compartments are reversible, requiring the input of ∼0.1 kJ/mol for approximately one minute to completely disassemble *in vitro* (Figure 2). This value provides a direct measurement of the energetic landscape that differentiates constitutive heterochromatin from euchromatin. Our experimental measurement is in remarkable agreement with that calculated using liquid Hi-C, a sequencing-based approach used to determine the critical length scale over which nuclear compartments are retained^72^. Using this method, Belaghzal, Borrman, and colleagues demonstrate that fragmentation of the genome to approximately 3 kbp resulted in nearly complete loss of A/B compartmentalization, however little information could be gleaned regarding the constitutive heterochromatin compartment due to the inability to assess contact frequencies for regions of repetitive C-Het DNA. The *in vitro* system characterized here provides direct quantitation of C-Het domain stability and places the complete energetic landscape that underlies nuclear compartmentalization between 0.1-0.5 kJ/mol-Kbp. This range provides a valuable frame of reference for the energy necessary to modulate, modify, or disrupt the network of chromatin contacts within which a wide array of nuclear bodies exist.

Along with the energy required to disrupt these heterochromatin domains, the reformation process could also be followed in our *in vitro* experiments. Almost immediately upon cooling back to room temperature, internal chromatin micro-domains start to form and eventually coalesce into macroscopic domains of similar morphology to those observed prior to heating (Figure 2B). During reformation, local regions of H3K9me3 enrichment are observed, which grow in a networked manner as more Me-chromatin is partitioned into the domain(s).

These disassembly and reassembly dynamics are akin to what takes place during initial establishment of C-Het domains in early Drosophila embryos. During development, C-Het initially displays elongated and highly amorphous configurations during the initial stages of reformation follow nuclear division, due to the underlying chromatin polymer, but then coalesces into more spherical compartments ^9,60^. In the *in vitro* system used here, the underlying chromatin is significantly shorter than that found *in vivo*, however elongated structures are also observed, suggesting a universal pathway of initial network formation followed by coalescence and relaxation to minimize the surface area of the Me- and U-chromatin compartments exposed to both solvent and each other.

### HP1a modulates chromatin solvation

The proclivity to minimize domain surface area is driven by the differential interaction parameters between the Me- and U-chromatin and the surrounding buffer, and the interaction between Me-chromatin and U-chromatin. We observe that HP1a modulates all of these interactions. In the absence of HP1a, both chromatin-types phase separate into homogeneously mixed condensates, indicating that Me- and U-chromatin interactions are approximately equal to the interaction of each chromatin type and the surrounding solvent. This provides evidence that H3K9me3 is not sufficient to drive macroscopic chromatin compartmentalization. To form co-existing immiscible domains requires that Me- and U-chromatin interactions are no longer equivalent. In the presence of HP1a, the observation of these discrete domains indicates that HP1a modulates Me- and U-chromatin interaction parameters. Furthermore, that these domains co-exist within a single condensate indicates a hierarchy of chromatin-solvent interactions. By assessing the surface area of the interfaces relative to the surface area of each compartment type, it is clear that U-chromatin domains are more often encapsulated by Me-chromatin domains (Figure 5C, SI Figure I, SI Figure 11, SI Figure 13). This implies that, in the presence of HP1a, the interfacial tension for U-chromatin and solvent is greater than that of Me-chromatin and solvent, thus the total energy of the system is reduced when the interface between U-chromatin and the surrounding buffer is minimized. This difference between chromatin-solvent interactions gives rise to the propensity of the Me-chromatin to wet along the U-chromatin domains, providing a physical explanation for the observed compartment morphologies. While there are certainly many other factors that position C-het at the nuclear periphery *in vivo*^1,12,13,73,74^, it is also interesting to consider the possibility that this configuration may also represent a global “energy minimum,” as it does for the *in vitro* system characterized here.

### HP1a increases chromatin liquidity

This hierarchy of interfacial tensions is informative when Me- or U-chromatin condensates are characterized separately in the presence of HP1a. It is not trivial to extract the precise values of interfacial tensions and internal viscosity, though their relation to one another can be quantified by tracking condensate fusion dynamics. The fusion dynamics of chromatin alone (no HP1a) indicate that Me-chromatin does not behave as a classic viscoelastic fluid, but U-chromatin does, having an inverse capillary velocity of ∼4.5 μm/s. HP1a promotes fusion events for both types of chromatin (Figure 4) and confers distinct viscoelastic character to each chromatin type; with a higher inverse capillary velocity (i.e. ratio of viscosity to surface tension) for Me-chromatin (η/γ =∼8.66 μm/s) than U-chromatin (η/γ =∼1.98 μm/s) (Figure 4D). When the chromatin types are mixed, the interfacial tension of U-chromatin can be inferred to be higher than that of Me-chromatin. This inequity allows the qualitative inference that HP1a increases the viscosity of Me-chromatin with respect to U-chromatin.

HP1a modulates both the viscosity and interfacial tensions of Me- and U-chromatin indicating a global role for HP1a in maintaining proper chromatin “solubility” in the nucleus. Indeed, HP1a is observed throughout the nucleus and has been shown to interact with euchromatic regions devoid of H3K9-methylation^5,75,76,15,44^. Several experiments using HP1a null mutants or depletion methods result in loss of compartmentalization strength, over-condensed chromosomes, severe mitotic defects, and misshapen nuclei ^2,5,77^. Our *in vitro* data suggest that loss of HP1a results in chromatin compaction and disrupted compartment integrity due to the global loss of chromatin liquidity within the nucleus, likely leading to increased self-association of chromatin.

### H3K9me3 modifications liberates linker-DNA length while increasing three-dimensional compaction

The *in vitro* data from this study demonstrates that HP1a substantially differentiates the material properties of Me- and U-chromatin, and that H3K9me3 alone is also sufficient to change chromatin dynamics at both the nano- and mesoscale. H3K9me3 modifications limit the growth rate of chromatin condensates, and these condensates do not behave as a viscoelastic material in contrast to unmodified chromatin condensates (Figure 3). At the nanoscale, H3K9me3 releases an average of 10bp of linker DNA and promoted increased intra-array compaction compared to U-chromatin arrays (Figure 2). We posit that these nanoscale changes along the nucleosome array limits the mesoscopic size of methyl-condensates through two mechanisms likely working in tandem: (1) shifting nucleosome spacing away from the condensate-favoring 10n+5 bp and (2) limiting inter-array interactions necessary for internal rearrangements.

Within the nucleus, electron microscopy studies have shown that interphase chromatin is not a homogeneous, contiguous chromatin structure but rather exists as ‘clutches’ of ∼4-10 nucleosomes^49^. These clutches interdigitate with one another and likely represent the packing unit of chromatin within the nucleus. Our data suggest that H3K9me3 modifications promote 3-dimensional chromatin fiber compaction by shifting the balance between intra- and inter-array interactions. Detailed structural and theoretical studies would be particularly informative to understand how the binding of HP1a within the context of a chromatin condensate may modulate the inter-array compaction favored by H3K9me3.

### The interfacial tension of heterochromatin compartments is governed by a hierarchy of HP1a interactions

Our data indicate that HP1a acts to increase chromatin liquidity and H3K9me3 promotes 3-dimensional chromatin compaction. When HP1a, Me-chromatin, and U-chromatin coexist, distinct compartments are formed with discrete and stable interfaces. Both the interfacial tension and viscosity of liquid-like condensates are governed by a combination of homo- and hetero-typic interactions between chromatin types, HP1a, and the surrounding solvent. These interactions are formally known as Flory χ parameters ^23,71^. When two polymers, in this case Me-chromatin and U-chromatin, have approximately equal χ parameters with respect to the solvent, mixtures of the two will produce homogenous co-condensates. This is indeed what we observe in the absence of HP1a (Figure 1). It should be noted that H3K9me3 is sufficient to change the growth rates of Me-condensates, suggesting that this epigenetic mark does modulate the χ parameter, but in a subtle manner such that it is not sufficient to prevent mixing with unmodified chromatin. Structural studies will be necessary to determine if this subtle change to χ is sufficient to drive microphase separation between Me- and U-chromatin arrays within these mixed condensates.

The formation of macroscopic immiscible compartments in the presence of HP1a indicates that the χ parameters, for Me-chromatin and the surrounding solvent, and U-chromatin and the surrounding solvent, are no longer equal. The degree of internal compartmentalization thus provides a direct way to assess how HP1a modulates the various χ parameters of Me- and U-chromatin. Our HP1a mutant analysis indicates that both dimerization and H3K9me3 binding are required for the creation of a stable interface. Thus, both HP1a-HP1a interactions and the scaffolding of HP1a to H3K9me3 are necessary to establish the hierarchy of interfacial tensions (γ_u-sol_ ≥ γ_u-me_ > γ_me-sol_) deduced from the observation that U-chromatin compartments were consistently surrounded by Me-chromatin domains. When the network of non-specific HP1a interactions is weakened by the charge-depleted hinge mutant, Me-chromatin is still observed to coat U-chromatin domains. However, there is an increase in Me- and U-chromatin mixing across this interface, due to increased interactions between Me- and U-chromatin. Increasing the affinity of the chromodomain for H3K9me3 decreases the domain overlap and specifically decreases the surface area of Me-domains at an interface. Thus indicating that increased HP1a retention on Me-chromatin increases the self-interaction of Me-chromatin at the expense of Me- and U-chromatin interactions.

These observations lead us to a model where the specific binding of HP1a molecules to H3K9me3 tails leads to the scaffolding of a highly dynamic HP1a-shell around Me-chromatin arrays, which promotes increased association of Me-chromatin via local HP1a interactions and a decrease in Me- and U-chromatin interactions. How sharp these interfaces are depends on the magnitude of HP1a-HP1a interactions and HP1a-H3K9me3 binding (Figure 6). HP1a plays a global role in defining the material properties of both U-chromatin and Me-chromatin, such that both compartments maintain liquidity and are capable of spontaneously compartmentalizing.

This suggests a global role for HP1a in modulating chromatin-solvent interaction parameters while also conferring distinct material properties to chromatin compartments in an H3K9me3-dependent manner. Our data quantify the energetic landscape of C-het compartments and suggest that HP1a may modulate how proteins interact with the underlying chromatin beyond a simple steric hindrance model, perhaps by directly modulating the intrinsic energetic landscape of chromatin domains at the level of chromatin solvation. The interfaces between Me-chromatin and U-chromatin created by HP1a likely represents a distinct chemical and biophysical environment within which biochemical processes, such as protein searches and enzyme activity, may take on unique characteristics.

## Author contributions

LDB designed, implemented, and analyzed the *in vitro* experiments with guidance from GK. HKK and JKR collected and analyzed the AFM data. SC performed the *in vivo* experiments. TE purified histones. The manuscript was written by LDB and GK with editorial feedback from HKK, JKR, SC, and SS.

## Acknowledgements

Sincere gratitude for all the members of the Karpen lab, current and past, for the feedback as this project evolved. Thank you to Professor Daniel Jost at the University of Lyon for encouraging the early stages of this work. Thank you to Professor Sam Safran for his enthusiasm for the project and input on the physics of phase-separating systems. Enormous thank you to Dr. Madeline Keenen (Duke University) who fundamentally collaborated on the initial conceptualization of this project and to Dr. Emily Wong (UCSF) for her knowledge and generosity during the early stages of this work.

## Supplemental Figure legends

**SI Figure 1:** HP1a-chromatin co-condensate morphologies Full field of view images for the HP1a-chromatin mixtures shown in Figure 5. Segmented domains illustrate the boundaries of each chromatin-domain. Scale bars are 10 μm.

**SI Movie 1-4:** 3-dimensional images of HP1a-chromatin co-condensate

In all movies, Me-chromatin is red and U-chromatin is cyan. HP1a is green. Scale bars are 0.5μm.

“231028_sidebyside-reslice_wt-HP1a” shows a side-by-side image stack of the same droplet shown along the X-Z (left) and Y-Z (right) planes.

“231028_zstack-wt-HP1a” is the same droplet as above, shown as a z-stack along the X-Y plane. A composite of the chromatin channels is shown on the left and HP1a is shown on the right.

“240125_wt_zstack” displays two droplets as a z-stack along the X-Y plane. “240125_wt-reslice” displays the same droplet(s) as an image stack along the X-Z plane.

**SI Figure 2:** Droplet fusions (A) Static images taken 15 minutes after Me-chromatin and U-chromatin were mixed. Composite images display droplets in various stages of mixing. Scale bar is 10μm. (B) Example time courses for HP1a-chromatin mixtures undergoing fusions. Time courses boxed in blue indicate fusions between U-chromatin domains. Time courses boxed in red show fusions between Me-chromatin domains. Time courses boxed in purple show fusion between Me- and U-chromatin domains. All images are 5 seconds apart, and all scale bars are 5μm.

**SI Movie 5:** chromatin-droplet fusion

“Combine_no-HP1a_me3-unmod-fusion” shows the Me-chromatin and U-chromatin droplet fusion without HP1a from Figure 1D. From left to right: Me-chromatin in red, U-chromatin in cyan, overlay. Frame rate is 10sec.

SI Movie 6: HP1a-chromatin co-condensate fusion

“Combine_crop2_wt-HP1a_fusion” shows the Me-chromatin and U-chromatin droplet fusion with HP1a from Figure 1D. On the left if the composite of Me-chromatin in red, U-chromatin in cyan, and HP1a is shown in the right. Frame rate is 2.5 sec.

**SI Figure 3:** Dual-color chromatin mixtures with HP1a (A) U-chromatin labeled with either cy3 (red) or cy5 (cyan) was mixed in the presence of 25 μM HP1a and the composite droplets were imaged 2 hours after mixing. (B) The same experimental set up as in (A) but with Me-chromatin. All scale bars are 10 μm

**SI Figure 4: *In vitro* and *in vivo* heating** (A) U- and Me-chromatin condensates were heated to the specified temperature, incubated for 5 minutes then imaged. (B) S2 cells expressing mGFP-HP1a and mScarlet H2B were heated from room temperature (22°C) to 37°C, incubated for 5 minutes, imaged, and then the heating turned off. Images of the same cells were captured every 15 minutes. Scale bars are 10 μm.

**SI Movie 7:** HP1a-chromatin co-condensate heating “230802_heatramp-realtime_combined_5fps” shows the movie for the data displayed in Figure 2A. From left to right: HP1a, Me-chromatin, U-chromatin. Frame rate is 5 frames per second.

**SI Figure 5:** Dextran partitioning into chromatin condensates (A) U-chromatin (cyan) and Me-chromatin (red) phase separation in the presence of 1mg/ml 500 kDa Dextran-fluorescein. Images were taken 2 hours after initial mixing. (B) U-chromatin (cyan) and Me-chromatin (red) phase separation in the presence of 1mg/ml 3 kDa Dextran-fluorescein. Images were taken 2 hours after initial mixing. Scale bars are 10 μm.

**SI Figure 6:** chromatin condensate material properties (A) Plots of fusion time-2 vs. average initial start diameter for Me-chromatin (red) or U-chromatin (cyan) in the absence (left) and presence (right) of 25 μM HP1a. Same data shown in Figure 3D and 4F, respectively, but with fit lines for the data in the absence of HP1a. (B) Side-by-side box plots for each fusion time component defined in the text for Me-chromatin or U-chromatin in the absence and presence of HP1a.

**SI Figure 7:** AFM methods for counting NCP occupancy and linker length **(A)** and **(B),** Representative AFM images of the crosslinked unmodified arrays and tri-methylated arrays (*n* = 8 independent experiments for both and *n* = 91 and 94 for unmodified arrays and tri-methylated arrays, respectively.). **(C),** The volume distribution of the Nucleosome Core Particles (NCPs) from crosslinked arrays. The volume of each masked NCP was measured, and then the volume of a single NCP is defined by the center value of the Gauss fit. (*n* = 44 and 53 nucleosomes for unmodified and tri-methylated arrays, respectively) **(D)** and (**E),** The number of the NCPs in the unmodified and in the tri-methylated arrays. We counted the number by measuring the volume of each nucleosomal array divided by the volume of a single NCP (See Method) (*n* = 44 and 53 nucleosomes for unmodified and tri-methylated arrays, respectively). **(F),** Representative image of DNA linkers of nucleosomal arrays. The curvy linker length was measured (red lines). (**G)**, Box plot of the linker length in the unmodified arrays and tri-methylated arrays (the box plots span from mean -s.d. to mean +s.d., the center thick lines show the mean, and the whiskers represent minimum and maximum values unless otherwise specified.) (*n* = 91 and 94 nucleosomal arrays and *n* = 494 and 540 linkers for unmodified and tri-methylated arrays, respectively).

**SI Figure 8:** AFM methods for measuring R_g_ **(A)** and **(B),** Representative AFM images of the unmodified arrays and the tri-methylated arrays (*n* = 2 independents experiments for both). (**C),** Schematic of the radius of gyration (*R_G_*) calculation. First, a nucleosomal array was cropped and masked to cover entire nucleosomes and DNA. To reduce the error, the background and the other particles were removed, and the center of the image and *R_G_*was calculated (See materials and methods for details). (**D),** Raw data of the *R_G_* versus volume scatter plot of unmodified arrays (*n* = 133 nucleosomal arrays). The nucleosomal arrays could be grouped into the three categories: (i) an oligomer of nucleosomal arrays, (ii) single nucleosome, and (iii) a dissociated nucleosomal array (right down panel). (**E**). Filtered data of the *R_G_* versus volume scatter plot of unmodified arrays (*n* = 96 nucleosomal arrays). To exactly analyze a single nucleosomal array, we excluded the oligomers and dissociated arrays, data points were filtered within the mean ± 3. (**F) and (G),** Box plot of the volume (**F**) and *R_G_*(**G**) of the unmodified and tri-methylated arrays from the filtered dataset (*n* = 96 and 114 arrays for unmodified and tri-methylated arrays, respectively).

**SI Figure 9:** Effect of H3K9me3 in cis and trans on droplet size **(A)** Histogram of droplet areas (in μm) for chromatin assembled with a 1:1 mixture of H3K9me3-nucleosome and unmodified nucleosomes Inset image shows a representative image for these condensates. Average area is 7.11μm^2^. (B) Histogram of droplet areas (in μm) for chromatin condensates formed with a 1:1 mixture of U-chromatin and Me-chromatin, as in Figure 1A in the absence of HP1a. Inset image shows a representative image for these condensates. Average area is 7.47 μm^2^. (C) shows the independent channels for Me-chromatin (red) and U-chromatin (cyan) for the inset image in (B). All scale bars are 10 μm.

**SI Figure 10:** HP1a-dependent condensate formation HP1a (green) titration with either Me-chromatin (red) or U-chromatin (cyan) at “low salt buffer” where chromatin does not phase separate. Buffer conditions were adapted from Keenen et al and are as follows: 25 mM Hepes pH 8, 75 mM KOAc, 1mM MgOAC, 1mM DTT, 1mg/ml BSA, 4% glycerol. Scale bars are 10 μm

**SI Figure 11:** In vivo HP1a-mutant imaging (A) Live-cell localization of meLAP-tagged HP1a mutants in S2 cells where endogenous HP1a has been RNAi depleted. Scale bar is 5 μm. (B) FRAP curves for HP1a constructs, measured on a DeltaVision microscope.

**SI Figure 12:** Mutant HP1a mixture component partitioning and shared interfaces (A) The same partitioning plot shown in Figure 5B, but for partitioning of the specified components within U-chromatin compartments. (B) Kernel Density Estimates for the fraction of Me-compartment perimeter (left plot) or U-compartment perimeter (right plot), at a shared interface, normalized to the total surface area of the “parent” droplet.

**SI Figure 13:** mutant HP1a-chromatin co-condensate morphologies Full field of view images for the HP1a-chromatin mixtures shown in Figure 5. Segmented domains illustrate the boundaries of each chromatin-domain. (A) AADA-HP1a mutant. (B) KED-HP1a mutant. Scale bars are 10 μm.

**SI Figure 14:** mutant HP1a-dependent condensate formation (A) KED-HP1a (green) titration with either Me-chromatin (red) or U-chromatin (cyan) at “low salt buffer” where chromatin does not phase separate. (B) I191E-HP1a at maximum concentration of protein achievable. (C) AADA-HP1a at maximum concentration of protein achievable. Buffer conditions were adapted from Keenen et al and are as follows: 25 mM Hepes pH 8, 75 mM KOAc, 1mM MgOAC, 1mM DTT, 1mg/ml BSA, 4% glycerol. Scale bars are 10 μm

## Methods

### Histone Purification

Plasmids for histone expression were a generous gift from Prof. Aaron Straight’s lab (Stanford) and were transformed into BL21 pLysS competent cells (NEB). 2-6 liters of E. coli were grown and induced according Dyer et al. Cell pellets were resuspended in 100 ml Buffer A (50mM Tris-HCl pH 7.5, 100mM NaCl, 1mM EDTA, 1mM Benzamidine, 2mM DTT) and sonicated on ice (1s on, 3 sec off at 70% power) for 10 minutes. The lysate was then centrifuged at 20,000g for 15 min at 4°C, the supernatant was discarded, and the pellet retained. Pellets were washed three times in 100 ml Buffer B (Buffer A + 1% Triton X-100) and centrifuged at 20,000g for 15 min.

Between each wash, the pellets were thoroughly broken up with a spatula. The pellets were then washed three times in 100 ml Buffer A, centrifuged at 20,000g for 15 min, with the pellets were broken up with a spatula between each wash. After washing, the inclusion bodies appeared uniformly white and flaky, an indicator of purity, and stored at −20°C.

For histone purification from inclusion bodies, 2 mL of DMSO was added to each inclusion body pellet and stirred at room temperature for 1 hour. This slurry was then dissolved in 50mL of Buffer C (6M Guanidinium-HCl, 50 mM Tris-HCl pH 7.5, 5mM BME), rocking, for 4 hr to overnight at room temperature. The mixture was then dialyzed in 6-8 kDa MWCO dialysis tubing against 2L of Buffer D (6M Urea, 20mM Tris-HCl pH7.5, 20mM NaCl; made fresh and deionized with MG AG 501-X8 (D) resin from Biorad). The dialysis buffer was changed 3 times, after 4 hours, overnight, and another 4 hours. The urea-dialyzed histones were spin down at 4000 x g, for 25 minutes at 4°C. The supernatant can be saved for classic ion exchange chromatography. The pellet was then resuspended in 50-100 mL Buffer C and rocked at room temp for 4 hours to overnight. This resuspension was spun down at 4000 x g, for 25 minutes at 4°C and the supernatant was dialyzed against 2L of MQ-H_2_O with 2mM DTT at 4°C. Three changes of the dialysis buffer were performed; after 4 hours, overnight, and a second 4 hours.

Protein concentration was measured as the absorbance at 276nm with a Nanodrop-One and aliquots were made for lyophilization. Proteins were lyophilized over 2-3 days to ensure complete dehydration and then stored at −80 until histone octamer refolding.

### Histone octamer refolding

Histone octamers were refolded according to Dyer et al 2003, using a 1.2:1.2:1:1 molar ratio of H2A:H2B:H3:H4. Refolded octamers were sized using an s200 size-exclusion column (Cytiva 17-5175-01), the octamer and dimer peaks were concentrated to >125 μM, at stored at −80°C until use.

### HP1a purification

6x-HIS tagged HP1 proteins were cloned into a pBH4 expression vector^1^, a generous gift from Dr. Coral Zhou, and then transformed into Rosetta competent cells (Millipore Sigma 70954). Protein expression was performed as published in Keenen et al 2020. Briefly, cells were grown at 37 C to an OD600 ∼1.0 in 1 L of 2xLB supplemented with 25 mg/mL chloramphenicol and 50 mg/mL carbenicillin. HP1 protein expression was induced by the addition of 0.4 mM isopropy-bD-thiogalactopyranoside (IPTG) and grown for an additional 3 hr at 37C, before pelleting at 4000xg for 30 min. Cell pellets were then resuspended in 30 mL Lysis Buffer (20 mM HEPES pH7.5, 300 mM NaCl, 10% glycerol, 7.5 mM Imidazole,1 mM PMSF (Milli-pore Sigma 78830). Cells were sonicated on ice (1s on, 3 sec off at 70% power) for 10 minutes.

Lysate was clarified by centrifugation at 25,000xg for 30 min. The supernatant was then added to 1 mL of Talon cobalt resin (Takara 635652) and incubated with rotation for 1 hr at 4 C, washed in a gravity column with 50 mL of Lysis Buffer, and eluted in 10 mL of elution buffer (20 mM HEPES pH 7.5, 150 mM KCl, 400 mM Imidazole). TEV protease (QB3, Berkeley) was added to cleave off the 6x-HIS tag and the protein mixture was dialyzed overnight in TEV cleavage buffer (20 mM HEPES pH 7.5, 150 mM KCl, 3 mM DTT) at 4 C. The cleaved protein was loaded on a Hi-Trap Q-column (Cytiva) and eluted by salt gradient from 150 mM to 800 mM KCl in buffer containing 20 mM HEPES pH 7.5 and 1 mM DTT. Protein containing fractions were collected and concentrated in a 10K spin concentrator (Amicon Z740171) to 0.2-1μM, 10% glycerol was added, and the protein was stored at −80 until use.

### DNA purification and labeling

The 12×194bp widom-601 array construct was generously provided by the Prof. Mike Rosen’s Lab at UT Southwestern^2^. Large scale purification was performed using the Qiagen Giga-prep kit. Briefly, DH 5-α (homemade) were freshly transformed with the 12×194-plasmid and selected using Carbenicillin. A single colony was picked the following day and used to inoculate 5mL of 2xLB, grown 8-10 hours at 37°C, and then transferred to 50 mL of 2xLB to grow overnight at 37°C. The overnight culture was then split across 5 L of 2xLB and grown 8-10 hours at 37°C. The culture was then pelleted, the pellet resuspended in Qiagen Giga-prep kit lysis buffer, and frozen at −20°C. These pellets were processed according to the Qiagen Giga-prep kit.

Once purified, the plasmid was digested using 20 U EcoRV (NEB) per mg of DNA at 37°C, overnight. The EcoRV was heat-inactivated per NEB instructions and the DNA was end labelled with Klenow polymerase (M0210S, NEB) using either Cy5-dCTP or Cy3-dUTP. Labelled DNA was ethanol precipitated and resuspended in 1xTE to a minimum concentration of 4mg/ml.

### Chromatin assembly

Nucleosome arrays were assembled using a standard salt dialysis method from Dyer et al^3^. The appropriate starting molar ratio of nucleosomes to 601-sequence was empirically determined for each histone octamer prep by preforming small scale assemblies and assessing 601-occupancy by digesting the array with EcoRI and running the mononucleosome products on a 4% native PAGE gel (cite). Assembly ratios ranged from 1.5-2.5:1, respectfully, with equimolar amounts of octamer and dimer added to each reaction. Once assembled, the 12 x nucleosome arrays were purified over a three part 8:18:25 w/v sucrose cushion. The sucrose cushions were centrifuged in a Beckman ultracentrifuge using a Ti-55 rotor, at 22,000 RPM, at 16°C, for 16 hours. Cushions were fractionated and fractions containing the assembled array were pooled and concentrated down using a 10K centrifugal filter (Ammincon Ultra, 0.5m, Millipore UFC501024). Arrays were concentrated to >3μM nucleosomes and stored at 4°C for immediate use or at −80°C for up to 6 months.

### Surface passivation

Surface blocking was adapted from Gibson et al 2020. Briefly, 384-well, glass-bottom plates (Griener) were cleaned with 2% helmanex (15 minutes), rinsed with 3 volumes MQ-H_2_O, then 0.5M NaOH (15 minutes), and rinsed with 3 volumes MQ-H_2_O. The wells were incubated overnight (4-deg, light protected) with 20 mg/ml PEG-silane in 90% EtOH. The PEG-silane was removed, the wells were washed with 2 volumes of 90% EtOH, and allowed to dry (in dark and protected from dust) overnight. Blocked wells were sealed with foil, and the plates stored at room-temp until use. For use, well were washed 3 times with MQ-H_2_O and blocked with 100mg/ml BSA for at least 1 hour at room temp. Immediately before imaging, the BSA was removed, the wells washed with two volumes of water and one volume of 1x phasing buffer, and the reaction mixtures are added immediately to the well.

### Phase separation assays

All chromatin droplets were formed at 1uM of 601-sites in 1x Phasing buffer. Phasing buffer was made as a 10x stock, and diluted down to a final buffer conditions of 25 mM Hepes pH 8, 150 mM KOAc, 1 mM MgOAc, 1 mM DTT, 1mg/mL BSA, and 4% glycerol. The final amount of 10x Phasing Buffer added to the reactions accounted for the salt from the HP1, thus ensuring all reactions contained the same amount of salts. All reactions were allowed to settle in the well for at least 30 minutes −2 hours and then imaged. For all static timepoints, images were acquired 2 after reaction mixtures were added to the well. For dynamic fusion tracking, images were acquired approximately 15 minutes following addition of reactions to the well.

### Imaging

All fluorescent images were collected on a Zeiss 880 line-scanning confocal with a 63x oil immersion objective. Brightfield images were collected on a Nikon Ti2, using a 100x objective.

### AFM measurement

We prepared dry-AFM samples for both linker length and Rg measurements (Figure 3E, F) by incubating U-chromatin or Me-chromatin in B1 buffer (25 mM HEPES pH 8, 75 mM NaCl, 2 mM MgCl2, 1 mM DTT) at room temperature and depositing them onto Poly-L-Ornithine (0.00001%) treated mica surfaces that were cleanly cleaved. For the linker-length measurement, to avoid dissociating nucleosomes, we crosslinked the nucleosomes with DNA using paraformaldehyde, while for the Rg analysis in Figure 3F, non-crosslinked nucleosomes were used to reflect natural nucleosome-nucleosome interactions. After washing the sample-deposited mica surfaces with 3 mL Milli-Q water, we dried the surfaces by blowing N2 gas using a gas gun. AFM imaging was performed in air using Peak Force Tapping mode with a Bruker NanoScope 6 with a set point of 150 pN. We used ScanAsyst-Air-HR cantilevers (spring constant: k = 0.4 N/m, tip-apex radius = 2 nm). We imaged the samples in a 5 μm × 5 μm scan with 5008 × 5008 pixels. All measurements were performed at room temperature and 50% humidity.

### AFM image analysis

Radius of gyration (*R*_g_) was calculated for each molecule in the AFM images. First, we cropped a single chromatin within the Region of Interest (ROI). The size of ROIs varied from 150 150 nm^2^ to 300 300 nm^2^ depending on the size of the molecule. After masking molecules in the ROI, the background outside the masked regions was removed. The center of mass of molecules and *R*_g_ were calculated from the images of the masked region of ROI. We defined the center of mass of molecules, and *R*_g_ as follows

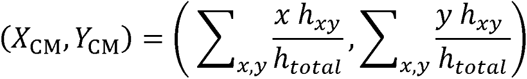

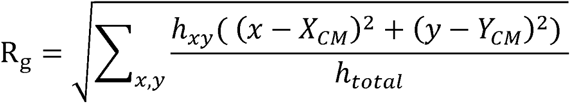

where *h_xy_* is the height at the pixel(*x, y*), and *h_total_* is the sum of *h_xy_* in the masked region.

### Image analysis

HP1a-and chromatin mixtures were analyzed using the Arivis software. Images of chromatin alone or single types of chromatin with HP1a were first processed in FIJI and then analyzed using a custom python segmentation code and downstream contour analysis. Code is available upon request.

**Droplet fusion analysis** was conducted in FIJI and the statistical analysis and plotting done in python.

### Partitioning Analysis

Single slices through the midplane of a field of droplets was selected for analysis in Arivis. The Me-chromatin and U-chromatin domains were each segmented using a simple thresholding method. These segments were then merged to represent the perimeter of the whole droplet. The total pixel intensity in the Me-chromatin, U-chromatin, and HP1a channels was measured with in each chromatin segment and for each “whole droplet.” Partitioning was then calculated as the total pixel intensity of a given channel in a specified segment divided by the total pixel intensity of that channel in the whole droplet (containing the specified segment).

### Shared Interface analysis

For each segment defined previously, the perimeter or area of each segment was also recorded along with the perimeter and area of the whole droplet within which segment was housed.

Overlap area between segments was calculated as the area shared between two segments and then normalized to the total area of their “parental” droplet. The quantification of shared interface area for either Me- and U-chromatin domains was calculated as the total perimeter of the interfaces within a single droplet normalized to the total perimeter of either Me- or U-chromatin domains.

### Contact Angle Analysis

Using a custom python code, available upon request, Me-chromatin compartments and U-chromatin compartments were segmented for a 2-D image through the midplane of a field of droplets. The contours for these segments were extracted using the skimage.measure.find_contours function. The intersections between contours were identified and at these points tangent lines were calculated for each contour and the angle between these intersecting tangents was calculated.

### Data analysis

Data visualization and analysis was performed using python. Code available upon request.

