## Supplementary material for "HP1a promotes chromatin liquidity and drives spontaneous heterochromatin compartmentalization": SI figures

**Supplemental Figure 1: HP1a - chromatin co-condensate morphologies**

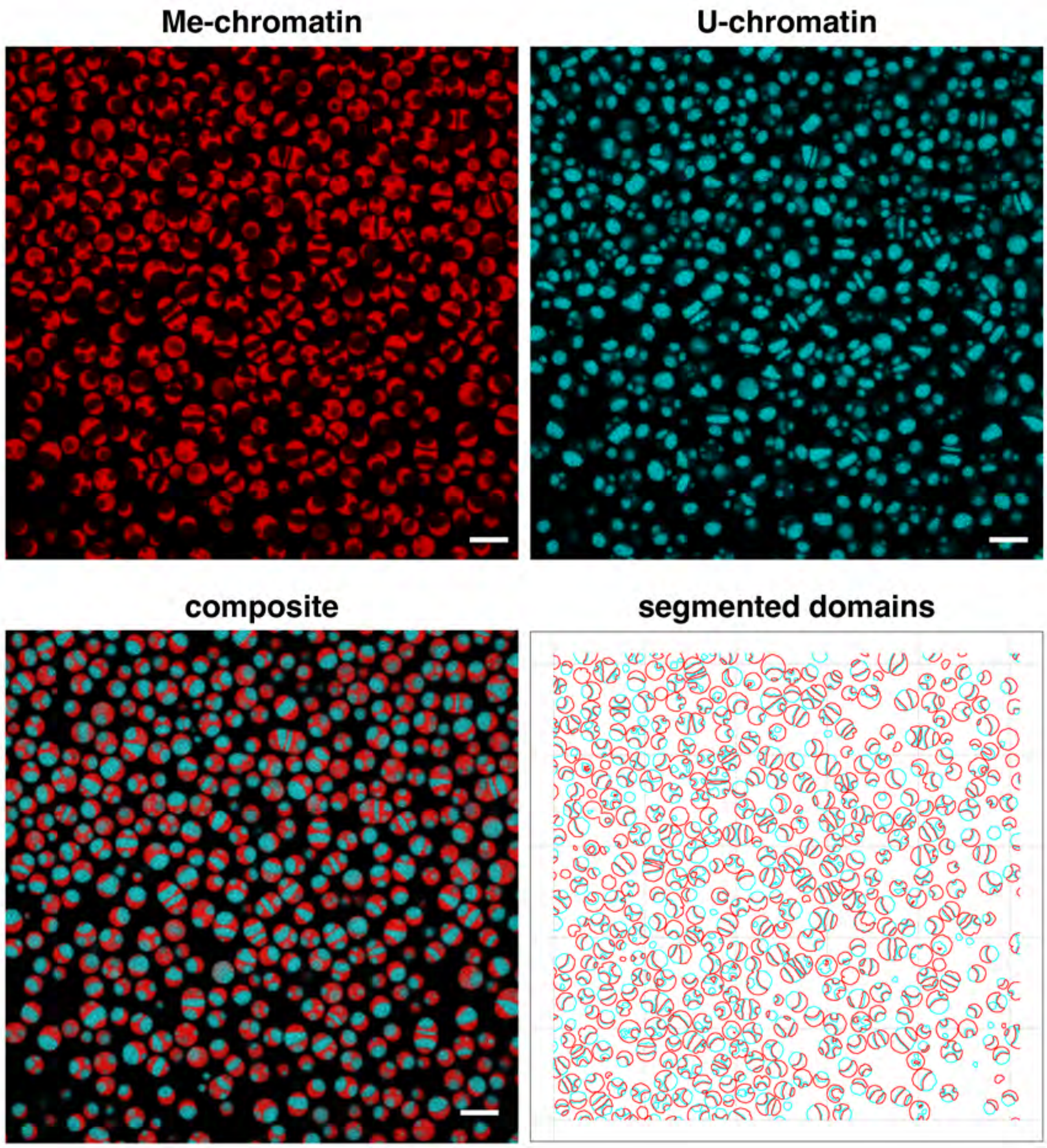

**Supplemental Figure 2: Droplet fusions**

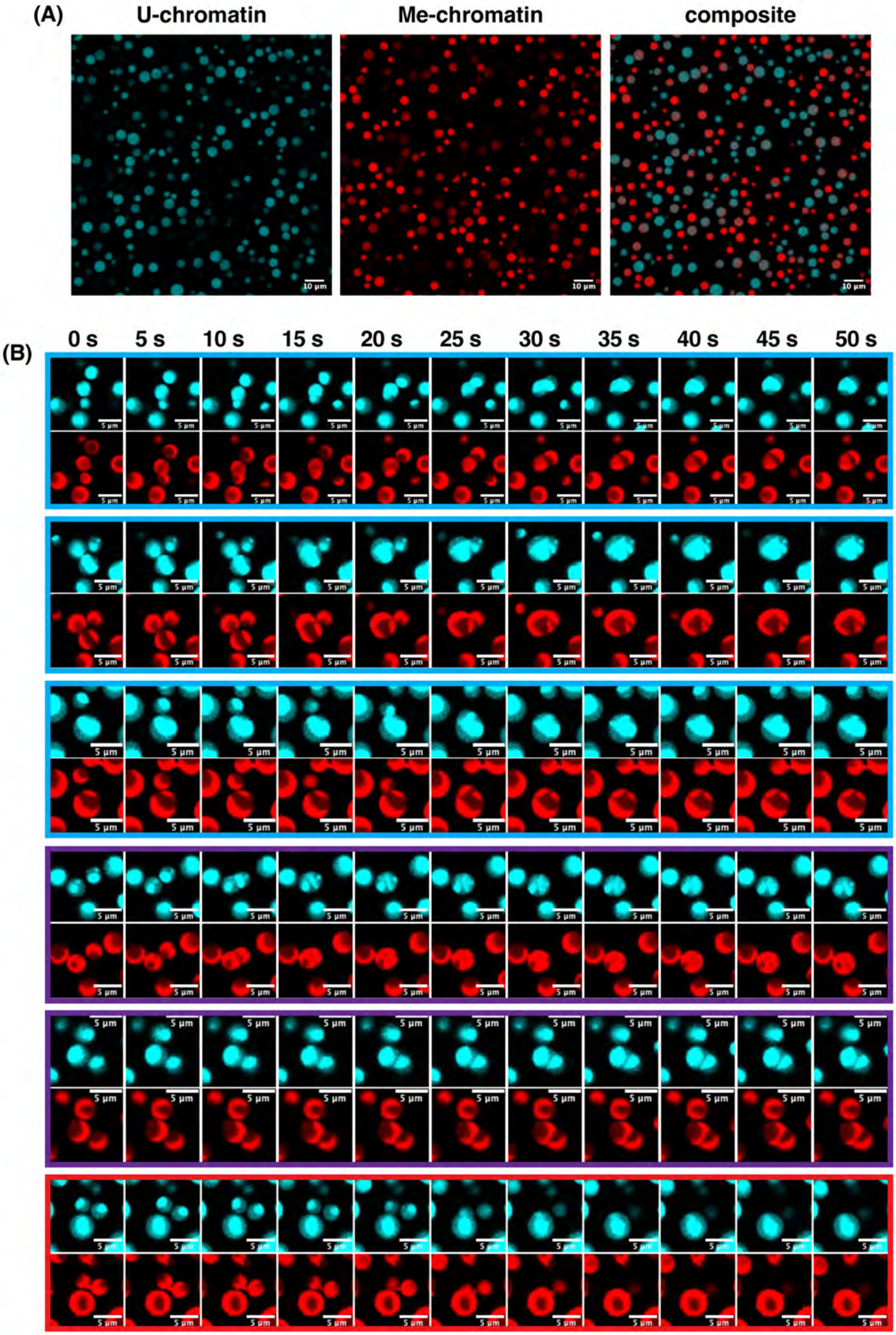

Supplemental Figure 3: Dual Color chromatin mixtures with HP1a

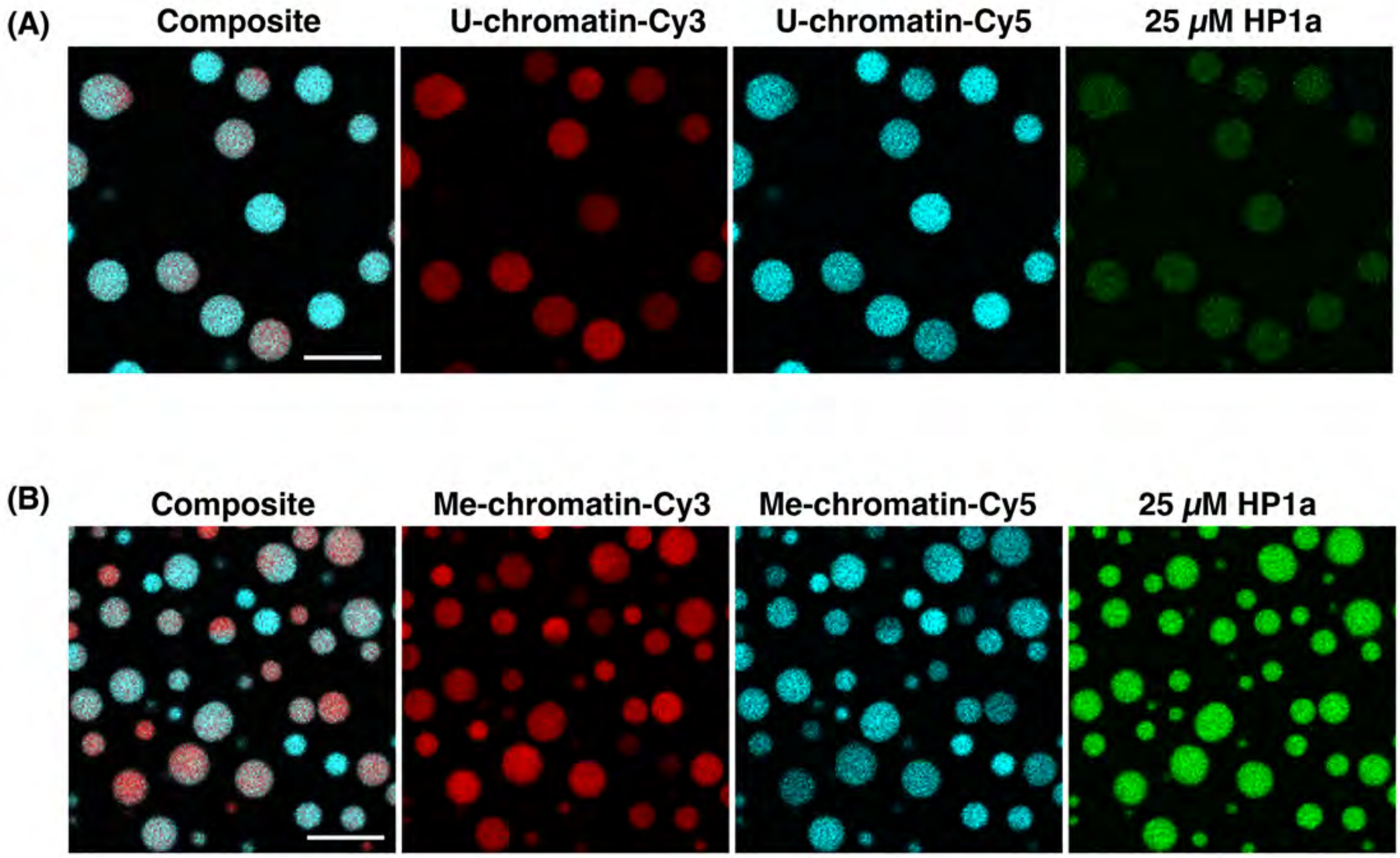

Supplemental Figure 4: heating experiments

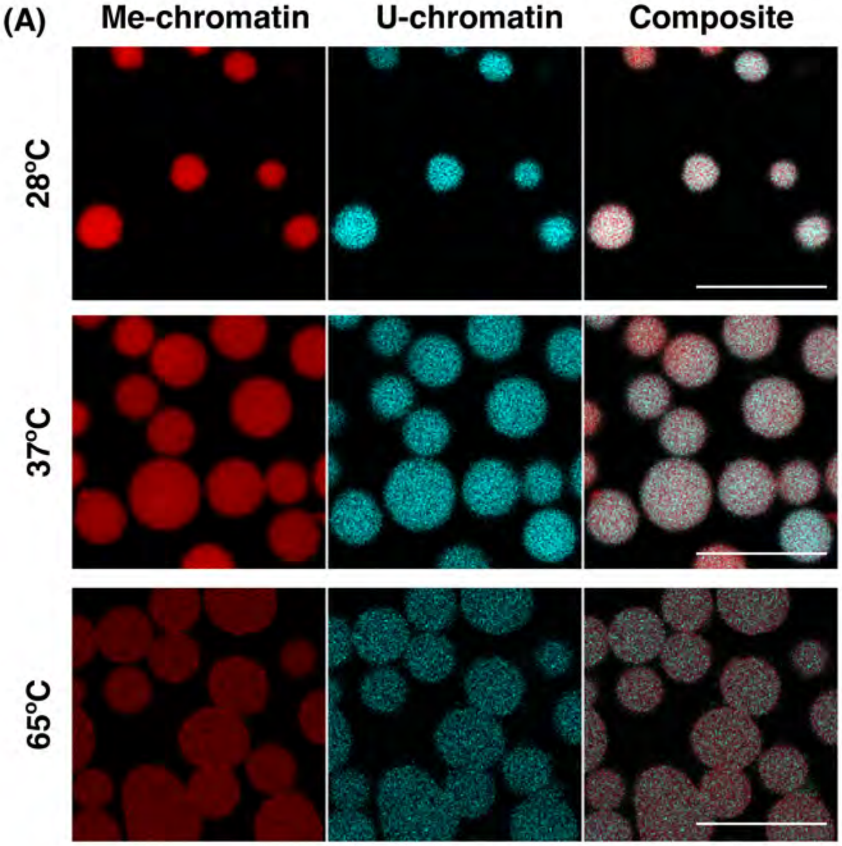


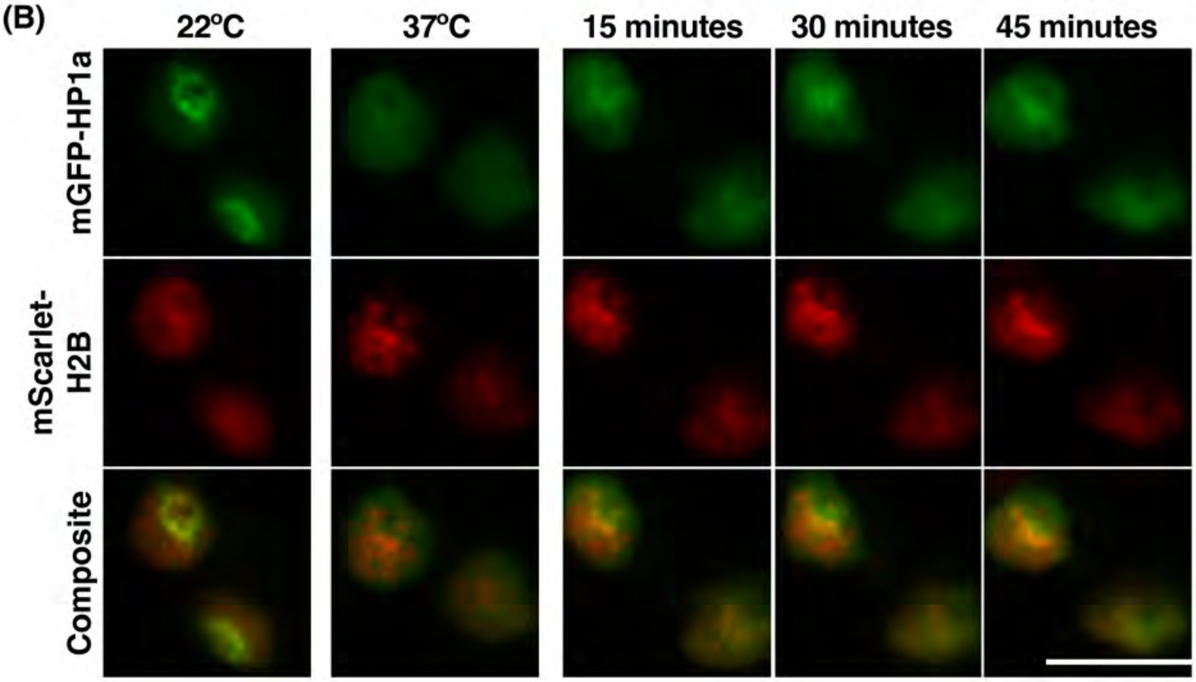

**Supplemental Figure 5: Dextran partitioning into chromatin condensates**

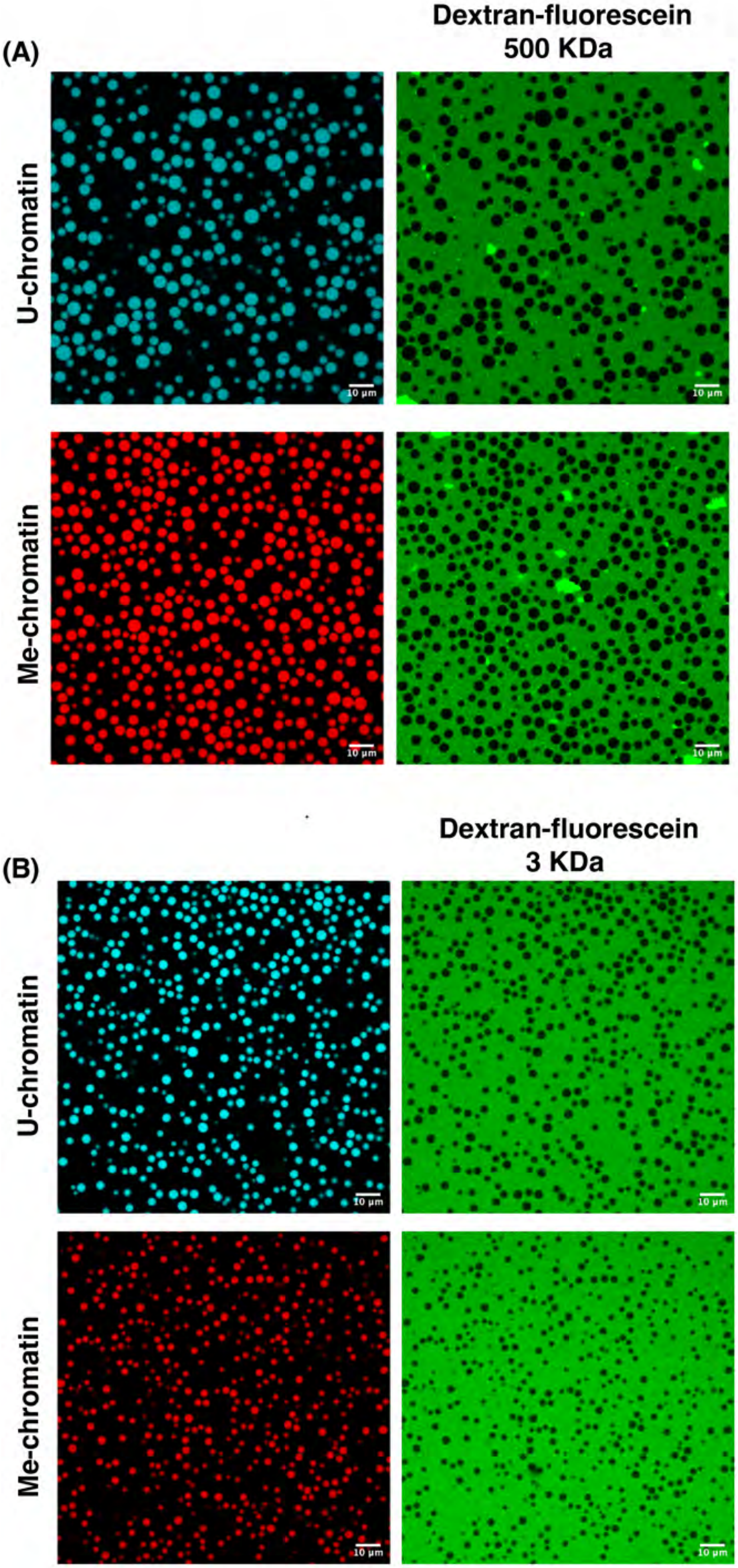

#### Supplemental Figure 6: Droplet fusion analysis side-by-side

(A)

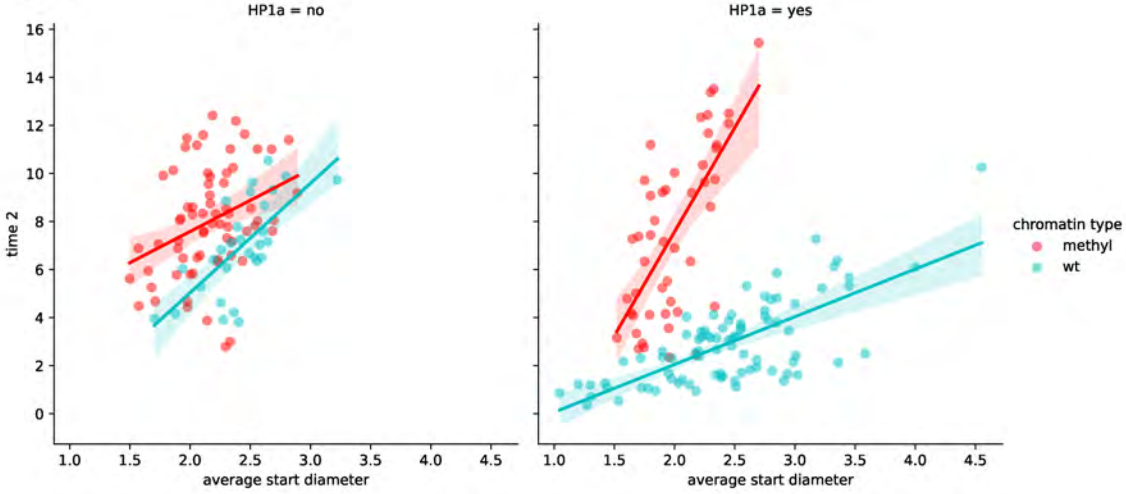


(B)

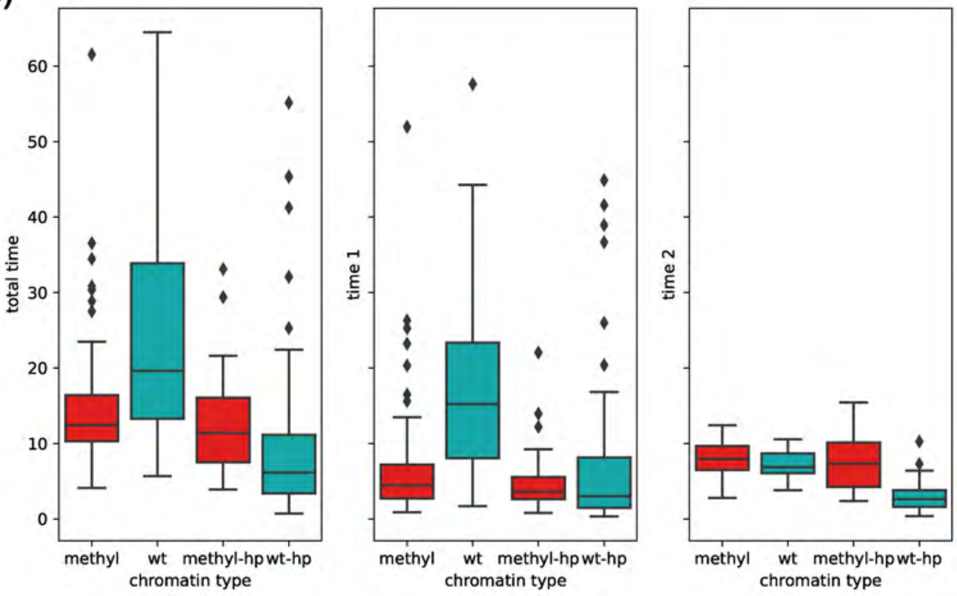

**Supplemental Figure 7: AFM methods for counting NCP occupancy and linker length**

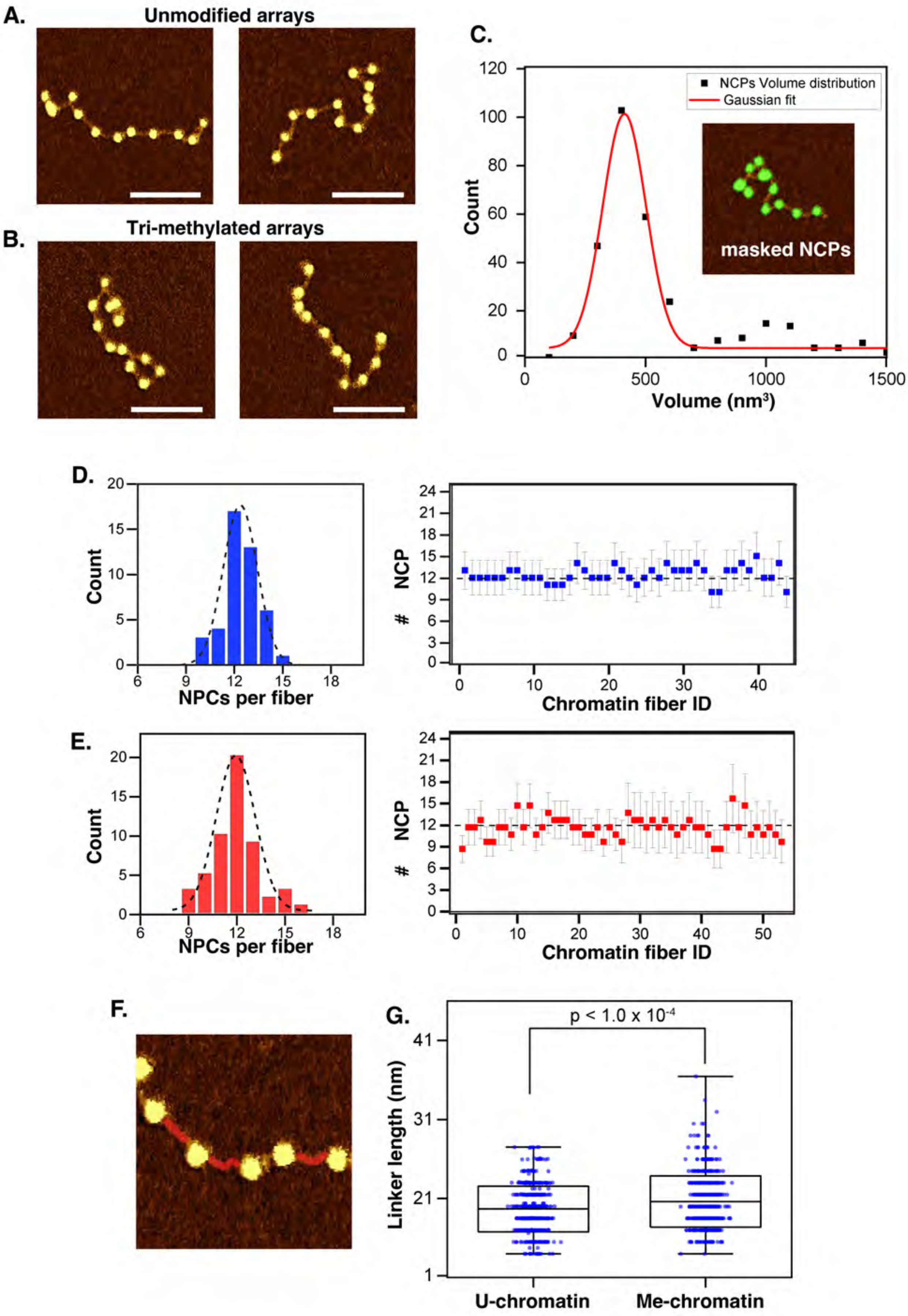

Supplemental FIgure 8: AFM methods for calculating $R_g$

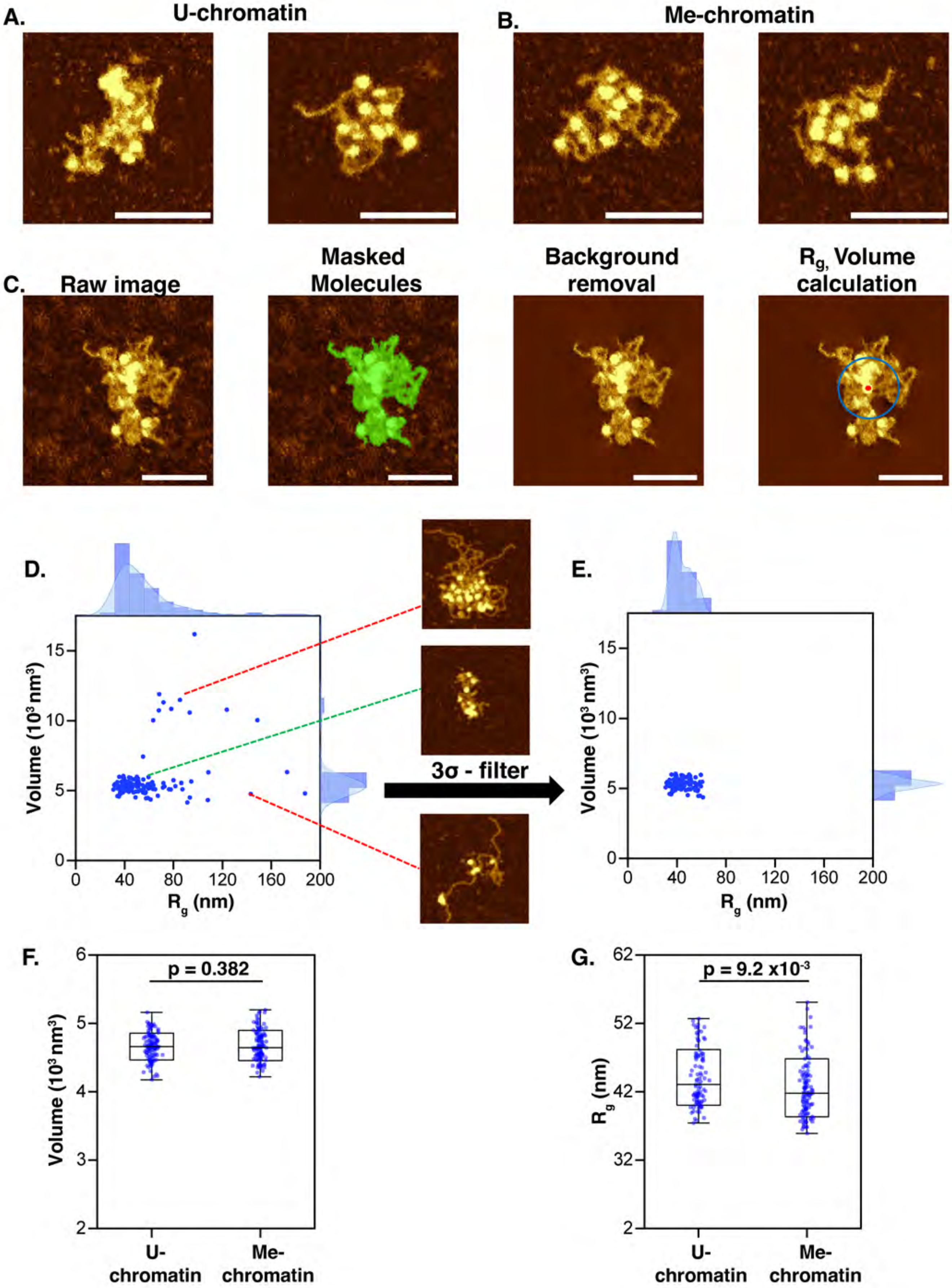

### Supplemental Figure 9: Effect of H3K9me3 in cis and trans on droplet size

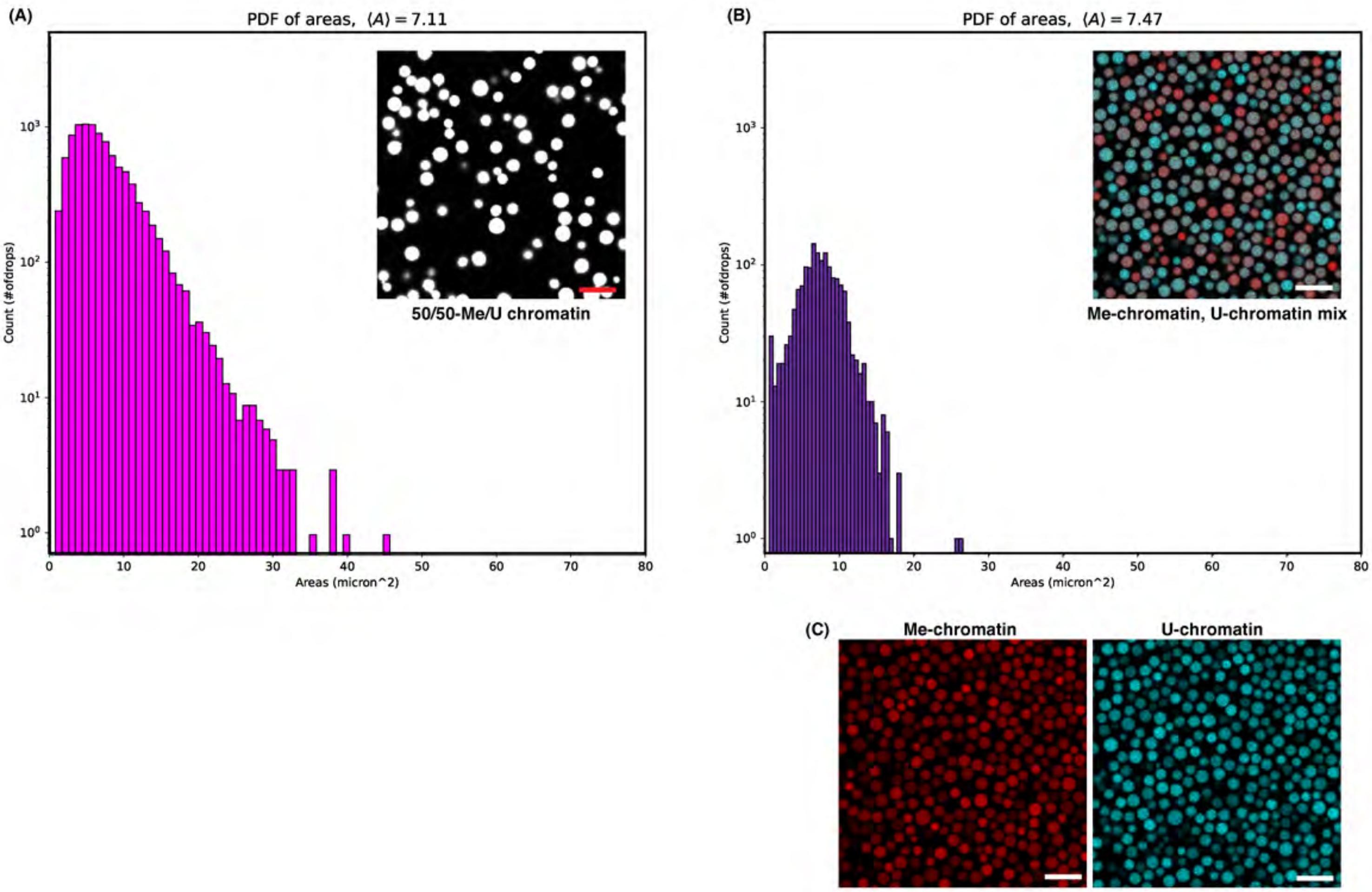

**Supplemental Figure 10: HP1a-dependent condensate formation**

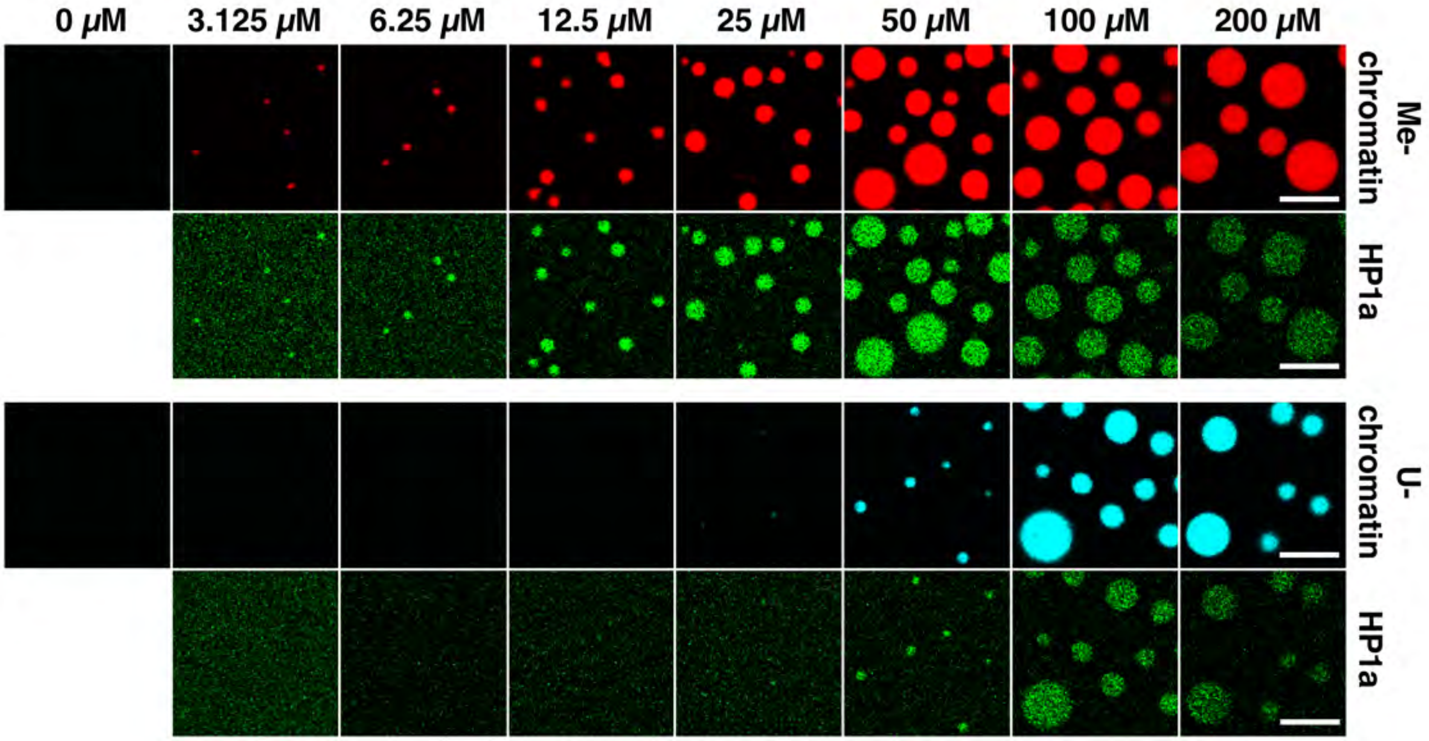

**SI Figure 11: In vivo imaging of HP1a mutants**

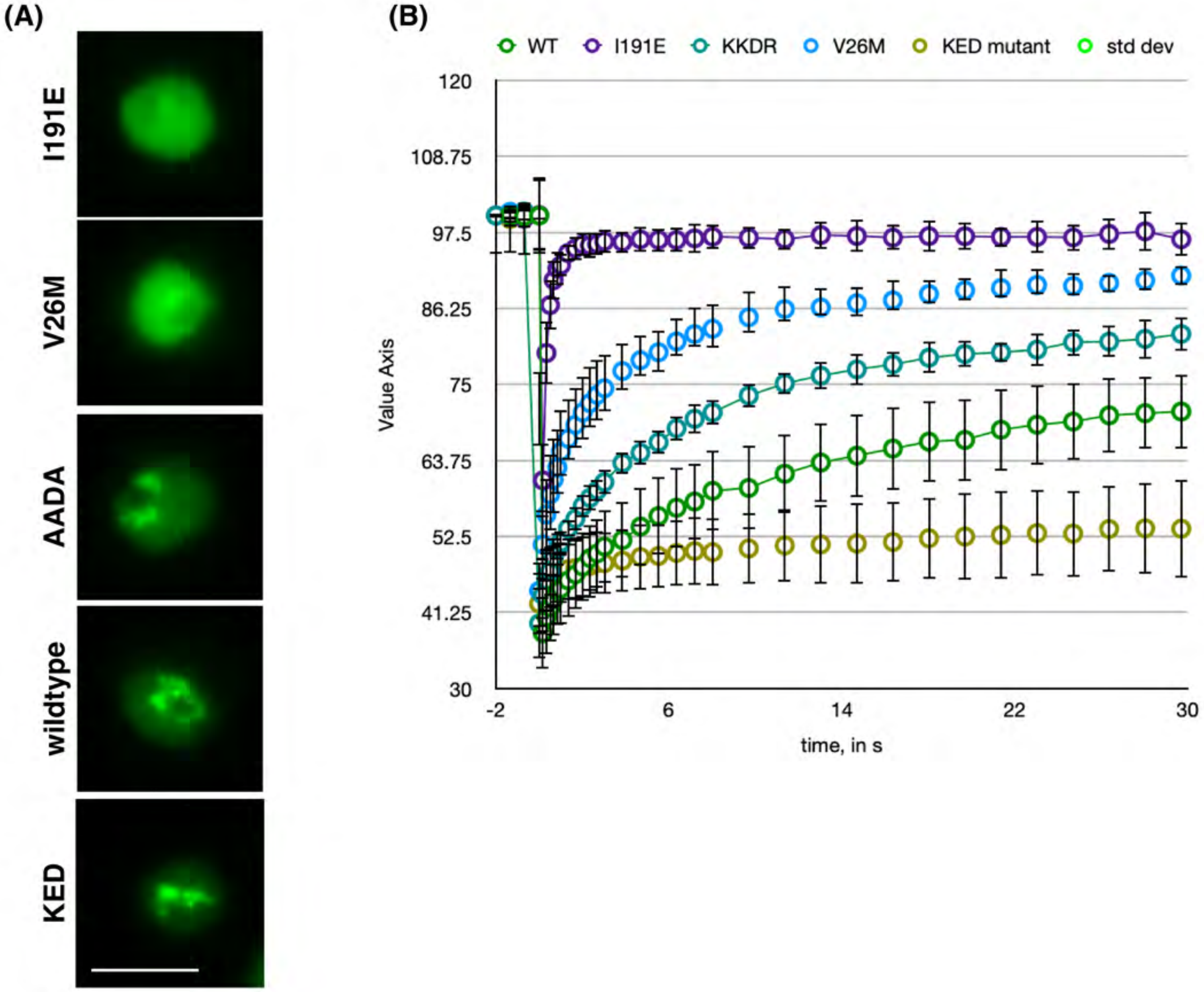

**Supplemental Figure 12: Mutant HP1a component partitioning and shared interfaces**

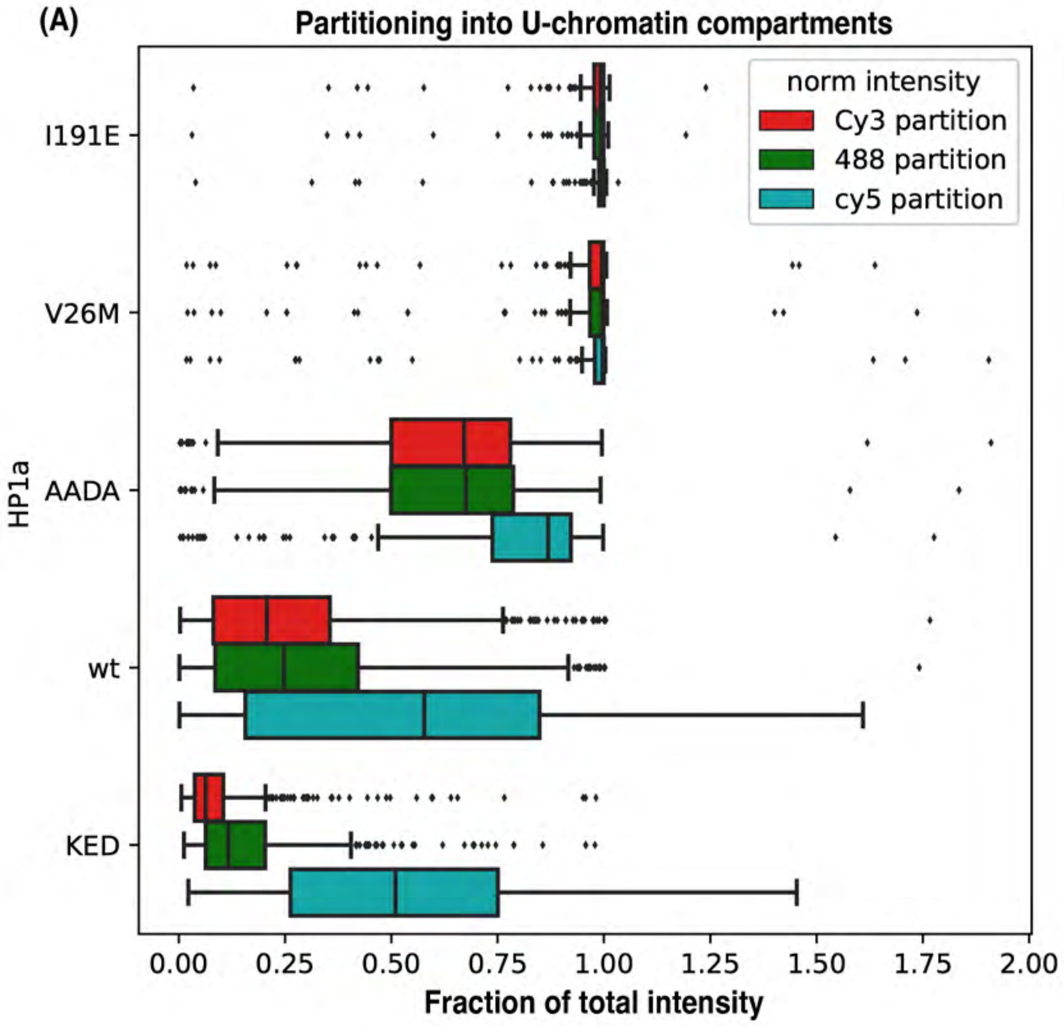


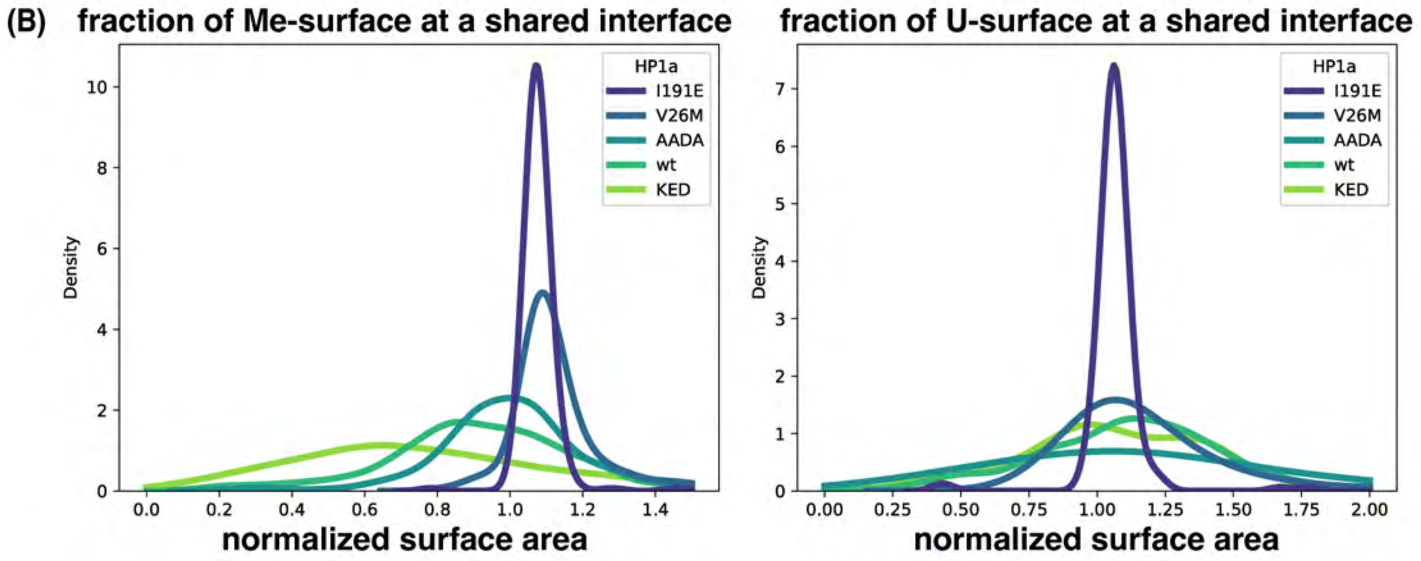

**Supplemental Figure 14: mutant HP1a-dependent condensate formation**

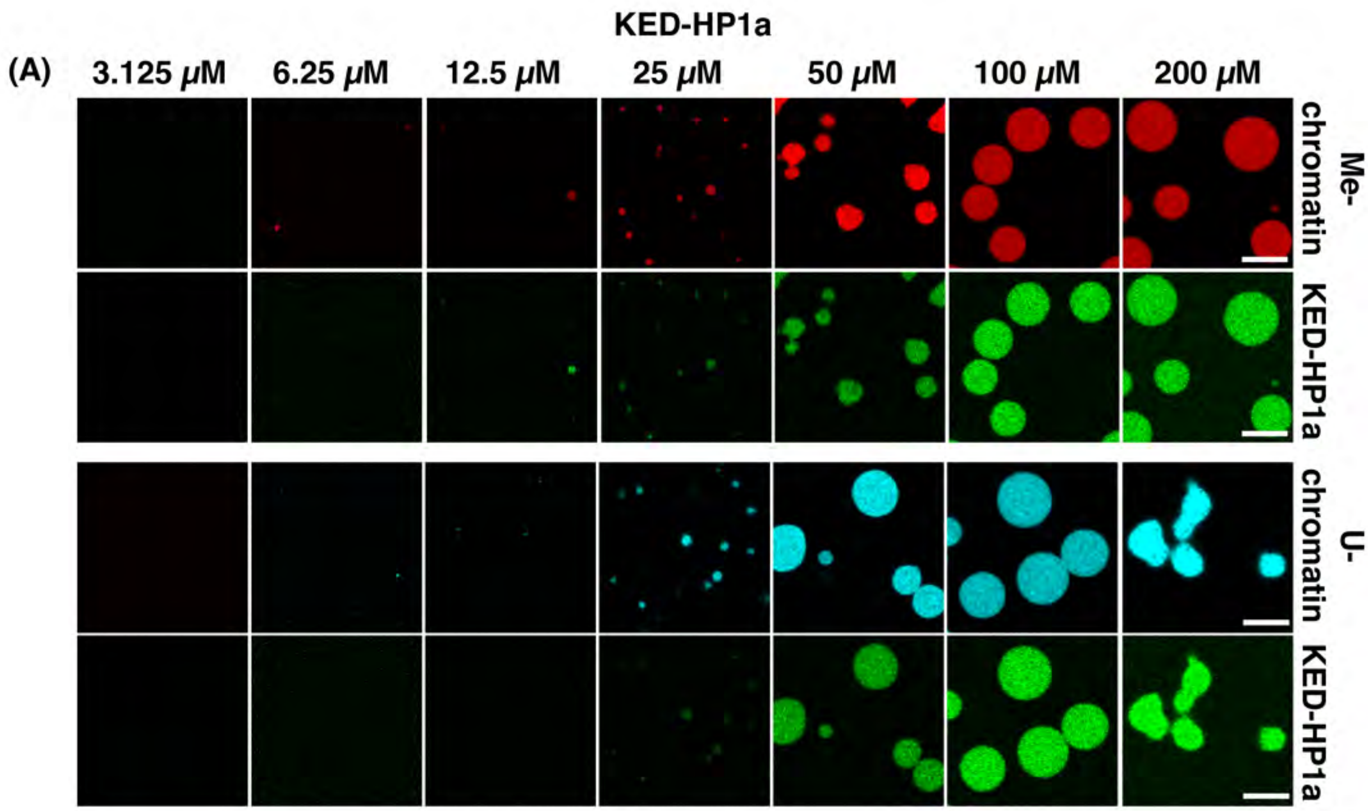


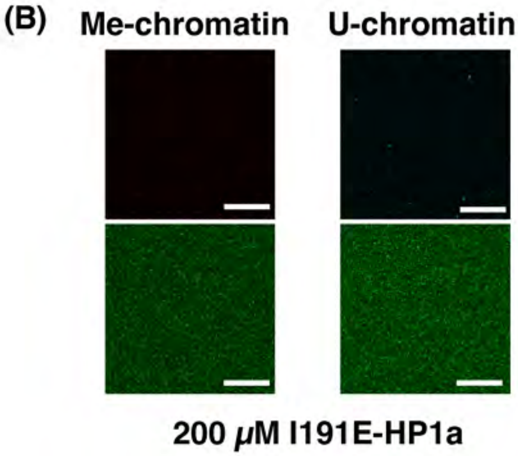


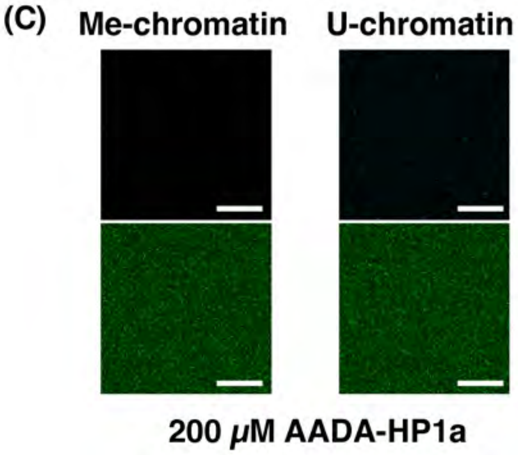

**Supplemental Figure 13: mutant HP1a-chromatin co-condensate morphologies**

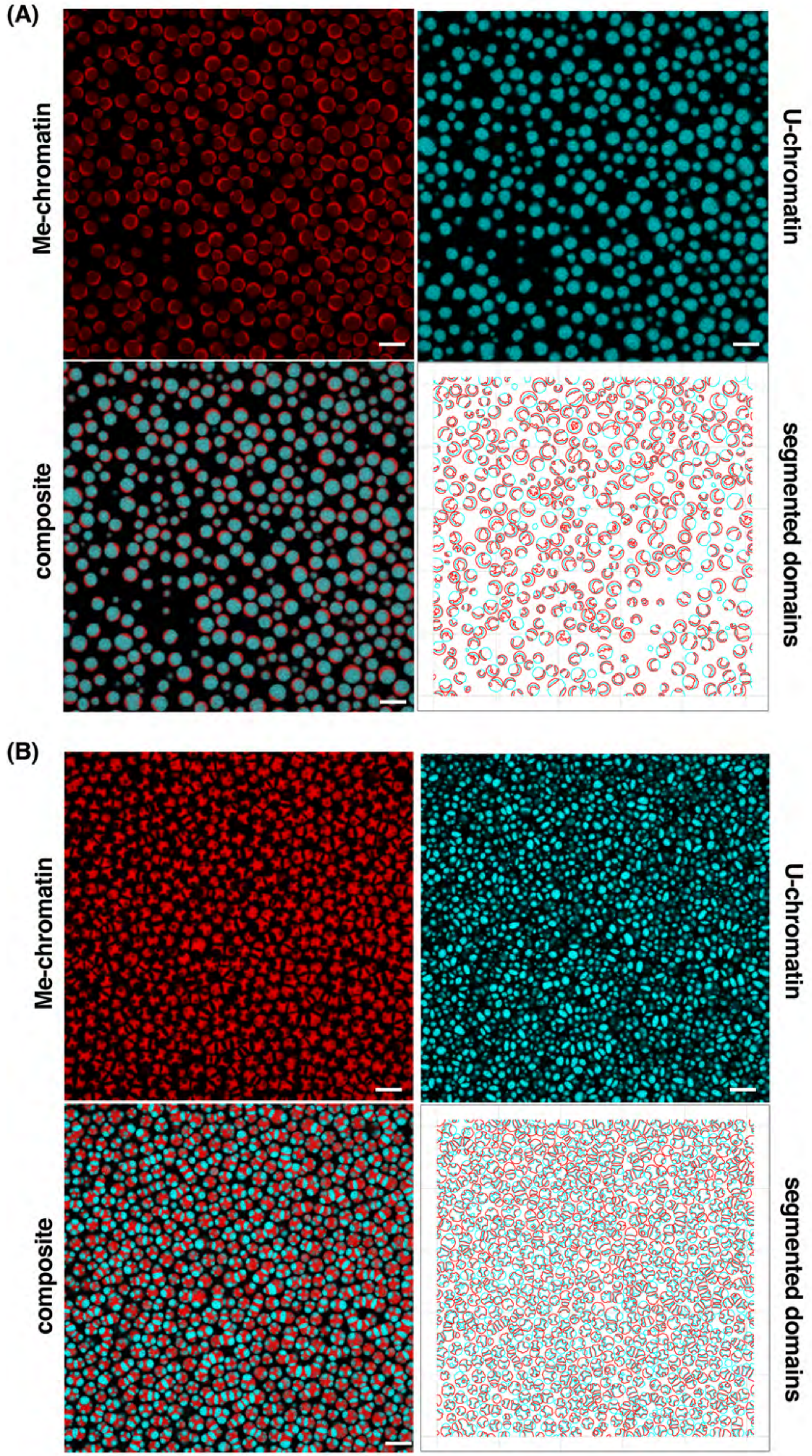
